# Anticipating xenogenic pollution at the source: Impact of sterilizations on DNA release from microbial cultures

**DOI:** 10.1101/833228

**Authors:** David Calderón Franco, Qingnan Lin, Mark C. M. van Loosdrecht, Ben Abbas, David G. Weissbrodt

**Author notes:** **Correspondence:** Prof. David Weissbrodt, Assistant Professor, Weissbrodt Group for Environmental Life Science Engineering, Environmental Biotechnology Section, Department of Biotechnology, Faculty of Applied Sciences, TU Delft, van der Maasweg 9, Building 58, 2629 HZ Delft, the Netherlands.

## Abstract

The dissemination of DNA and xenogenic elements across waterways is under scientific and public spotlight due to new gene-editing tools, such as do-it-yourself (DIY) CRISPR-Cas kits. Over decades, prevention of spread of genetically modified organisms (GMOs), antimicrobial resistances (AMR), and pathogens from transgenic systems has focused on microbial inactivation. However, sterilization methods have not been assessed for DNA release and integrity. Here, we investigated the fate of intracellular DNA from cultures of model prokaryotic (*Escherichia coli*) and eukaryotic (*Saccharomyces cerevisiae*) cells, commonly used as microbial hosts for genetic modifications, such as in white biotechnology. DNA release was tracked during exposure of these cultures to conventional sterilization methods. Autoclaving, disinfection with glutaraldehyde, and microwaving are used to inactivate broths, healthcare equipment, and GMOs produced at kitchen table. The results show that current sterilization methods are effective on microorganism inactivation but do not safeguard an aqueous residue exempt of biologically reusable xenogenic material, being regular autoclaving the most severe DNA-affecting method. Reappraisal of sterilization methods is required along with risk assessment on the emission of DNA fragments in urban systems and nature.

**Graphical abstract:** 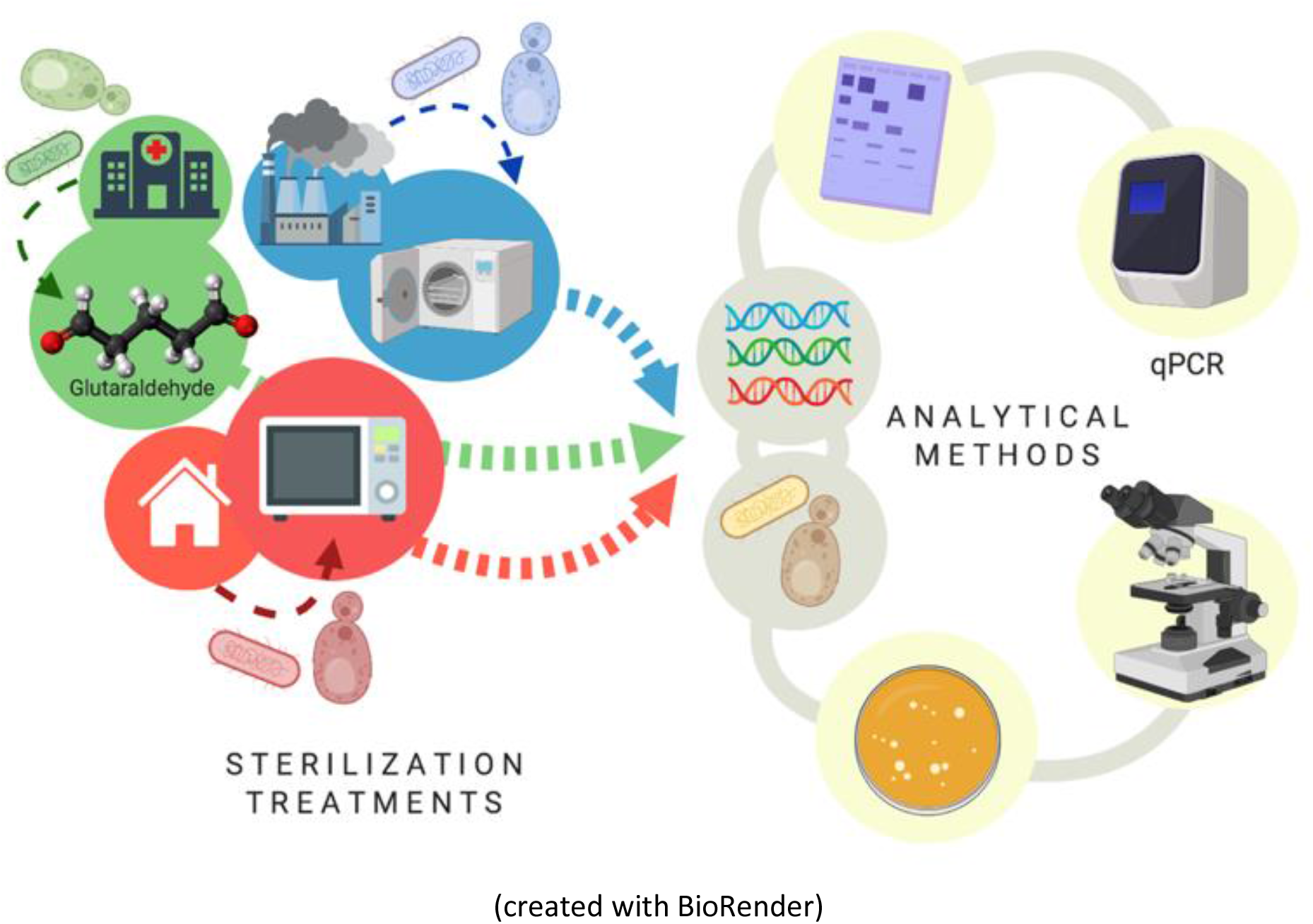

## Introduction

The rapid development of gene-editing tools together with the broad applications of genetically modified organisms (GMOs) have triggered biosafety concern on the hazard composed by the dissemination of unwanted DNA into the environment after sterilization ^1^ Concerns have been risen on the emission of xenogenic and mobile genetic elements that may carry antibiotic resistance genes (ARGs) or pathogenicity, and their transport across urban waterways through wastewater treatment plants into nature ^2,3^. Novel CRISPR-Cas technologies propel the engineering of microorganisms out of industry boundary with do-it-yourself (DIY) kits available at kitchen table. Uncontrolled, diffuse emission of GMO materials via domestic waste streams could be a threat.

Current sterilization methods are proven to efficiently inactivate microorganisms. However, a key knowledge gap remains on their impact on DNA and its potential release into industrial, clinical, and domestic sewage. Common methods used to treat industrial broths, healthcare equipment and surfaces, and domestic waste primarily involve autoclaving, glutaraldehyde, and microwaving, respectively.

Several studies have shown how DNA present in food products react with different sterilization procedures ^4–6^. Treatments such as irradiation and autoclaving affect DNA in meat products or edible seeds by decreasing the total DNA content as well as causing DNA fragmentation, degradation and denaturation ^7,8^. However, the impact highly depends on the cell type, the sterilization method, and the process conditions.

Temperature, pressure, pH, and sterilization times significantly exert effects on DNA quality. For instance, temperatures over 100 °C have resulted in significant DNA strand clipping and irreversible loss of secondary structure ^4^. Normal autoclaving (121°C between 5 to 20 min) of food and crops did not impeded it to be available for PCR amplification ^4,5,9^

Microwaving is commonly used in kitchen procedures such as water boiling and food heating ^10,11^. It has been suggested to effectively kill bacteria, yeast, and molds on kitchen sponges ^12^. Microwaves at frequencies of 2450 MHz have been used to sterilize soil due to its ability to inhibit nitrification and sulfur oxidations ^13–15^. It has also been used in laboratory settings for pharmaceutical glass vials, culture media, or clinical specimens sterilization ^16^. The thermal effect mechanism is based on the absorption of microwave heat energy by the cell constituents, which leads to fast vibrations of cell membrane lipids resulting in the emergence of pores ^17^. These pores may cause leakage of vital intracellular molecules being able to cause cell death^17^. High temperatures denature cellular biomolecules such as proteins, which may also be a reason of cells lysis ^18^.

Glutaraldehyde is commonly used in industry, research labs and, more specific, in hospitals as a disinfectant on dental and medical instruments, such as endoscopes, and surfaces. Glutaraldehyde has a wide range of biocidal activity against both Gram-positive and Gram-negative bacteria, viruses, and spores ^19,20^. It is a strong cross-linker that combines with multiple molecular functions such as amino and sulfhydryl groups ^21^. Glutaraldehyde affects cells by binding to nucleic acids and cross-linking enzymes responsible for oxygen uptake, destroying secondary structures and therefore causing disfunction of cytoplasmic molecules and death of cells ^22^.

Here, the model bacterium *Escherichia coli* and yeast *Saccharomyces cerevisiae* were used as model organisms to test the impact of autoclaving, microwaving, and glutaraldehyde on the release of DNA from microbial cultures of prokaryotes and eukaryotes, respectively. We elucidated the effects of these three dominant sterilization methods on DNA release, fragmentation, degradation, and amplification capacity on top of cellular inactivation, morphology, and integrity. This incepting work provides first insights to foster the management of the emission of xenogenic pollution.

## Results and discussion

### Sterilization methods inactivated over 99% of living bacterial and yeast cells

The microwave effect on cell viability (Fig. 1**a**) showed similar end-points but different profiles for both model prokaryotic and eukaryotic microorganisms. For both *E. coli* and *S. cerevisiae*, the microwaving performance for 15 s was sufficient to reduce the number of cells count to nearly 10% of control untreated samples, indicating severe inactivation effect on both microorganisms. Thermal effect is responsible for absorption of microwave energy by cell molecules, producing general heating of the cell ^23^. This causes disarrangement of cell membrane and disruption of cell wall structures by destroying the lipopolysaccharides and peptidoglycan of the cell surface. This results in the emergence of pores, cell aggregations, cytoplasmic proteins aggregation, and changes of membrane permeability ^17,24^, explaining why biomolecules such as proteins or nucleic acids ^15^ are detected in the extracellular fraction.

**Figure 1.**
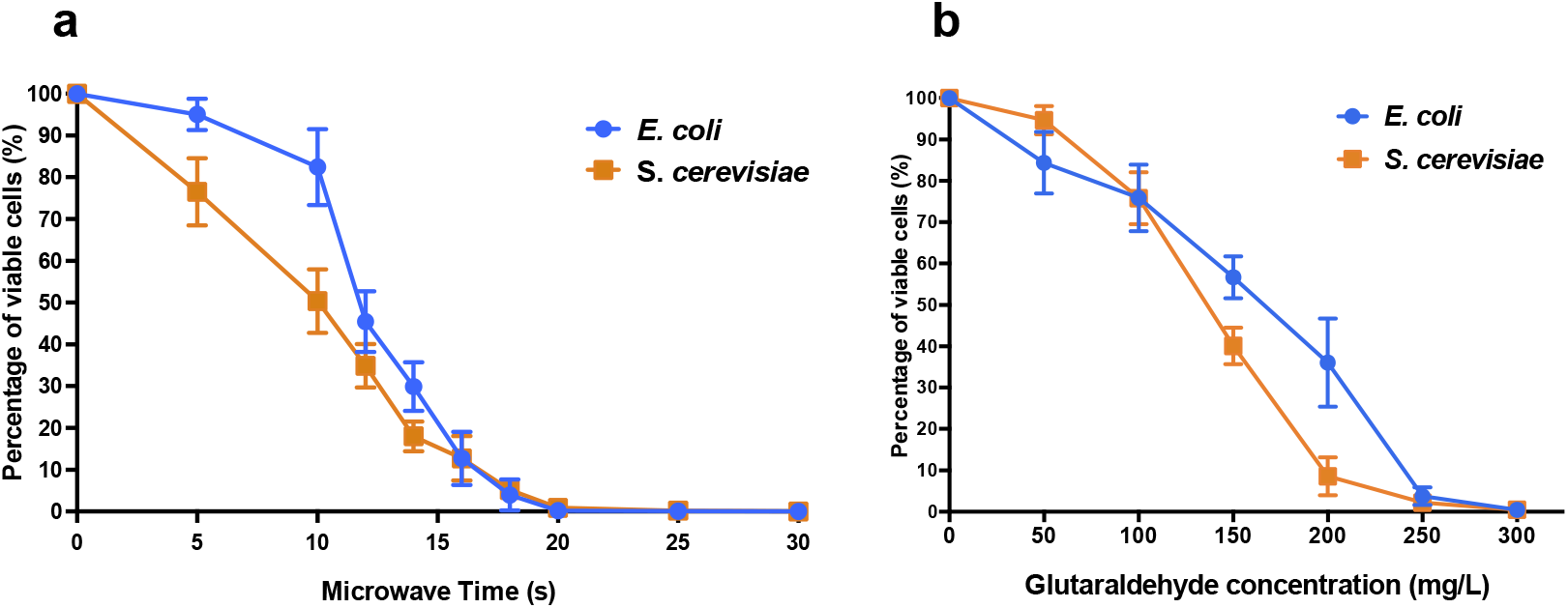
*E. coli* and *S. cerevisiae* cell viability after (**a**) different microwave exposure times and (**b**) glutaraldehyde concentrations. The percentage of viable cells is calculated against corresponding control sample cells (untreated with microwave and untreated with glutaraldehyde, respectively).

The biocidal effects of the increasing dose of glutaraldehyde were shown on both types of microorganism cells (Fig. 1**b**). *E. coli* and *S. cerevisiae* both displayed significant reduction of cell viabilities compared to their non-treated control samples. When 300 mg L^−1^ glutaraldehyde was applied, both microorganisms decreased to a range between 0.1% to 1% viable cells when compared to control cultures.

The autoclaving treatment was the most effective on both *E. coli* and *S. cerevisiae* (Table S1). All types of autoclaving program decreased the viable cell numbers under the minimum detection limit of <100 CFU mL^−1^, thus resulting in nearly zero survivor cells in the autoclaved cell suspension.

### Cell lysis and aggregation after autoclaving and microwaving whereas loss of cell transparency is common after all sterilization methods

Phase-contrast microscopy images showed that untreated *E. coli* cells were intact and displayed long rod-shaped structures with smooth surface (Fig. 2, left column). Microwave treatment after 10 s did not affect the overall *E. coli* cell structure as most of the cells maintained their shape (Supplementary Fig. 1). After 25 s, cells showed considerable cell debris as well as non-conventional shapes and cell aggregations. Cells transparency was also fully lost after 15 s, becoming dark non-conventional shaped cells compared to the non-treated cells. Short-term microwave treatment preserved cell structure (Supplementary Fig. 2) but displayed cell aggregations. The visible damage of *S. cerevisiae* cells emerged from 25 s, where cell walls destruction resulted in the fusion of *S. cerevisiae* cells into a pool of broken cells. Loss of cell transparency was also observed.

**Figure 2.**
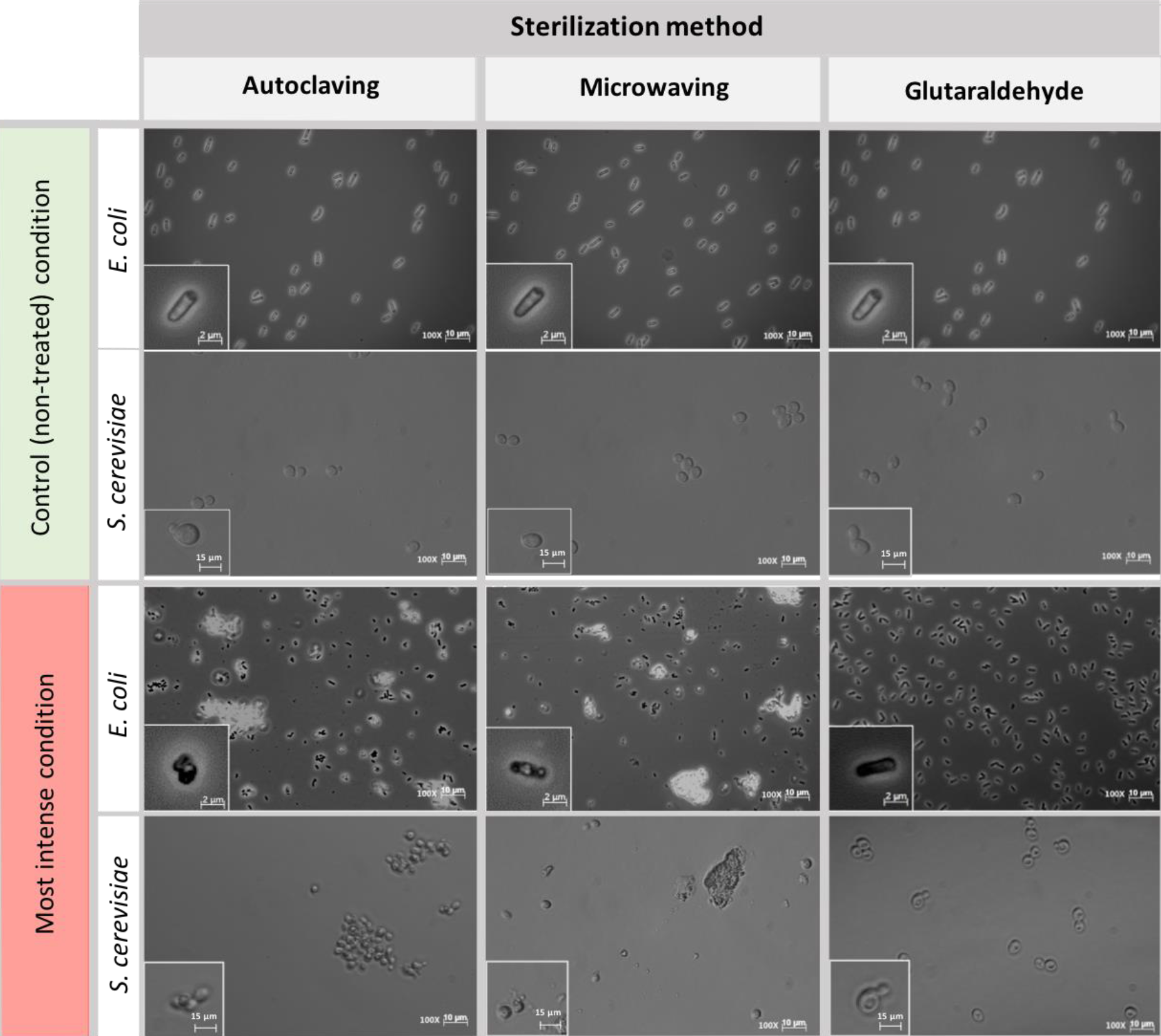
*E. coli* and *S. cerevisiae* cell morphology before and after being treated with the harshest sterilization method condition. Pictures were taken at 100x magnification. Control (non-treated) condition: microscopic pictures after overnight cultivation. **Left column:** Microwaving effect. Most intense condition corresponds to 30 s microwaves exposure time. **Middle column:** Autoclaving effect. Most intense condition corresponds to P4 (121 ^<u>°</u>^C – 30 min). **Right column**: Glutaraldehyde effect. Most intense condition corresponds to 300 mg L^−1^ glutaraldehyde. The morphological structure of single cells at each time point are shown at bottom left of each picture.

*E. coli* cells displayed severe structural damages under autoclaving treatment as cells no longer preserved the transparent rod-shaped morphology presented in control samples (Fig. 2, middle column). Cells were mostly ruptured into pieces of debris, twisted and shrunk, filled with denatured intracellular molecules (Supplementary Fig.3). For *S. cerevisiae*, under the first three types of autoclaving programs, cells maintained completely their spherical shape, and hardly any wreckages of cells were shown in suspension (Supplementary Fig. 4). Cell metamorphosis occurred under the highest intensity of autoclaving (Fig. 2, middle column) where cells lost their clear surface layer and started to perform partial fusions.

The most obvious effect of glutaraldehyde fixation on the morphology of both *E. coli* and *S. cerevisiae* cells (Fig. 2, right column) was the change of cell transparency. The intracellular area of *E. coli* cells turned black when fixed with glutaraldehyde and for *S. cerevisiae*, dark spots were present in their cytoplasm. No cell aggregation nor significant cell damage was observed. Cells from both microorganisms still preserved their structural frame even when the highest concentration of glutaraldehyde of 300 mg L^−1^ was applied (Supplementary Fig. 5 and 6).

### Significant DNA release observed under microwave and autoclaving sterilizations

The microwave effect on the total DNA released by *E. coli* was observed after 12 s and increased abruptly after 16 s of exposure (Fig. 3**a**). In *S. cerevisiae* cultures, the total DNA released increased constantly from the beginning of the treatment (Fig. 3**a**). Even if *S. cerevisiae* cells released DNA constantly from the start, their cell wall protective ability was higher than the *E. coli* ones: yeasts displayed higher resistance in terms of cell structural collapse and DNA release into the extracellular media. This links to the microscopic pictures presented in Fig. 2, where after 20 s under the microwaves, the *S. cerevisiae* cells preserved their structure whereas the *E. coli* cells lysed and aggregated (Supplementary Fig. **1** and **2**). Resistance differences also resulted in a lower percentage of total DNA released from *S. cerevisiae* at maximum exposure (30 s) of microwave sterilization. Nearly 35% of total DNA (73.9 ng μL^−1^) was released from *S. cerevisiae* cells at 30 s, in contrast to almost 50% (59.1 ng μL^−1^) from *E. coli* (Supplementary Fig. 7**a-d**).

**Figure 3.**
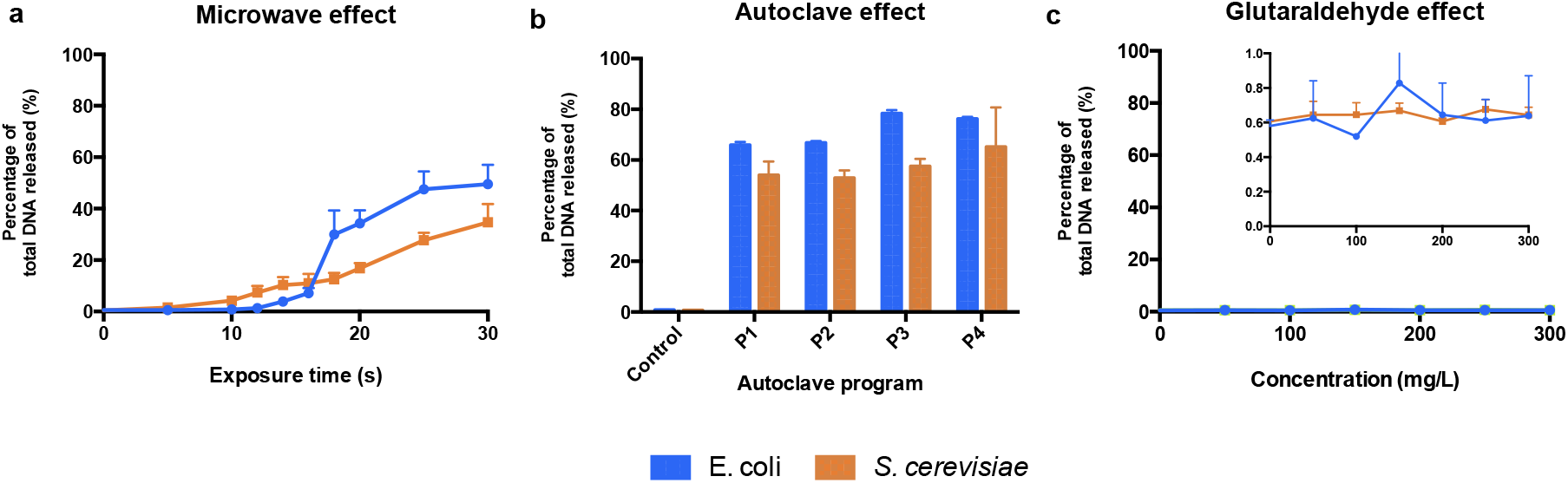
Total DNA released from *E. coli* and *S. cerevisiae* treated with (**a**) different microwave exposure times, (**b**) different autoclave programs and (**c**) different glutaraldehyde concentrations. The percentage shows the ratios of the amount of DNA (dsDNA) released in the supernatant against the total amount of DNA (released and remained combined). Autoclave programs: P1 (110 °C – 20 min), P2 (110 °C – 30 min), P3 (121 °C – 20 min) and P4 (121 °C – 30 min). Control (non-treated cells). 100% correspond to 59.1 ng μL^−1^ and 73.9 ng μL^−1^ for *E. coli* and *S. cerevisiae*, respectively.

All types of autoclaving led to considerable amount of DNA released to the medium from both microorganisms (Fig. 3**b**). Different autoclaving programs caused 65% (P1) to nearly 80% (P4) of total DNA leakage from *E. coli* cells while lower amounts of total DNA, 53% (P1) to 65% (P4), were observed on *S. cerevisiae* cultures (Supplementary Fig. **8**).

When an increasing amount of glutaraldehyde was applied from 0 to 300 mg L^−1^, the percentage of total DNA released to extracellular medium (Fig. 3**c**) stayed either steady at 0.6% for *S. cerevisiae* or fluctuating from 0.4 to 0.9 % for *E. coli*. Almost all the DNA from both microorganisms remained intracellularly (Supplementary Fig. 7**e-h**). Glutaraldehyde displays a protective effect against cell lysis ^25,26^ and strongly inhibits autolytic and proteolytic processes ^27^. This explains why DNA was not released into the extracellular fraction. Glutaraldehyde does perform cross-linking reactions with compounds present in the cell outer layers, thus negatively altering the permeability and transportability of cell membranes ^19,22,28^. Our observations show that this chemical exposure time (20 min) is enough to inactivate cells and prevent the transport and release of nucleic acids across membranes.

### Long microwave exposures and all the autoclave programs showed severe DNA damage

The DNA fragments of the untreated *E. coli* control sample displayed on agarose gel have a size above 10 kb. When cells were treated with increasing duration of microwave, the intracellular DNA bands were less intense and displayed a slight decline gradient of DNA sizes, trailed by smears at different degrees (Supplementary Fig. 9**a**). For *S. cerevisiae* (Supplementary Fig. 9**c-d**), the DNA extracted from untreated cells showed multiple bands of various lengths from approximately 400 bp to over 10 kb. After 25 s of microwaving, no clear bands but smears were displayed on the gels. An exposure of 30 s resulted in elimination of *S. cerevisiae* visible bands in the intracellular fraction (Supplementary Fig. 9**c**). The fragmentation patterns in *E. coli* matched the measurements of DNA content (Fig. 3**a**) for which a significant amount of DNA was released from 18 s onwards (Supplementary Fig. 9**b**). An increasing gradient of the free-floating DNA intensity on the gel from *S. cerevisiae* (Supplementary Fig. 9**d**) fitted to the gradual DNA release (Fig. 3**a**).

In contrast to microwaving, extracellular and intracellular DNAs from both microorganisms were more highly fragmented and/or degraded when heat and pressure (1.1 atm overpressure) were applied, displaying smears on agarose gels (Supplementary Fig. 10**a-d**). The highest autoclaving program (121 ^<u>°</u>^C, 30 min) showed the strongest DNA damage and release when compared with controls and other conditions (Supplementary Fig. 5**a-c**, lanes 5). Even when the harshest autoclaving was applied, intense bands of DNA were still observed on the agarose gels. Intracellular DNA degradation degree was lower than the released DNA after autoclaving treatment, presumably due to the protection of cells on its cytoplasmic DNA against external damage ^1,29,30^. A possible reason why *E. coli* shows higher resistance to stress when compared to *S. cerevisiae*, apart from their higher surface to volume ratio, could be its polyunsaturated fatty acids (PUFAs) composition in their membrane: they contribute to cell membrane flexibility. PUFAs level in *S. cerevisiae* membranes are low or inexistent when growing under normal conditions ^31,32^ whereas in *E. coli* cells are higher ^33^.

Intracellular DNA of *E. coli* treated with the highest concentration of glutaraldehyde resulted a smear with high fragment lengths (Supplementary Fig.11**a**, lane 7). Intracellular genomic DNA on *S. cerevisiae* did not result in the absence of DNA bands but a decrease of the band intensity (Supplementary Fig. 11**c**, lane 7). The extracellular DNA from both types of microorganisms (Fig. 11**b-d**) showed similar patterns containing short DNA fragments from <100 bp to >200 bp, same as their corresponding untreated controls. The presence of residual extracellular DNA primarily results from the natural DNA release of microbial cells even before application of glutaraldehyde ^34^.

### Autoclaving was the most effective in compromising PCR-ability

The amplification efficiency of selected genes was assessed after sterilization by qPCR. Differences of log_10_ number of DNA copies per mL between bead-milled control samples (autoclaving and microwaving) or non-bead-milled control samples (glutaraldehyde) and treated samples gave an insight on the DNA integrity after sterilization. For autoclaving and microwaving, log_10_ values close to the control indicated a high number of amplifiable DNA sequences available in the sample. This meant that DNA was not degraded enough and maintained its integrity. For glutaraldehyde, values close to the non-bead-milled samples indicated no DNA release nor an effect on PCR ability.

In *E. coli* cultures, no effect on PCR ability was observed when DNA was released from cells after microwaving. There were higher initial (5 s) log_10_ differences with the bead-milled control sample (0.77 ± 0.02) values due to lack of DNA available on the extracellular fraction (6.79 log_10_ gene copies mL^−1^). DNA was exponentially released after 25 s (7.22 ± 0.07 log_10_ gene copies mL^−1^, Fig. 4**a**), and significantly released after 70 s, 7.84 ± 0.01 log_10_ gene copies mL^−1^). After 100 s exposure, DNA was released from cells (7.81 ± 0.08 log_10_ DNA copies) as its number of sequences even got higher than the control values (7.56 ± 0.01 log_10_ DNA copies) but its PCR ability was not compromised. Regarding autoclaving, a signal was observed even after P3 and P4 programs were applied (Fig. 4**c**): 1.28 ± 0.11 and 1.16 ± 0.04 log_10_ gene copies per mL difference with the bead-milled control, respectively. Glutaraldehyde did not have a significant effect on the PCR ability mainly because samples treated with glutaraldehyde did not release DNA (Fig. 4**e**). An analysis of variance (ANOVA) on these scores yielded significant variation among autoclaving and microwaving treatments but not glutaraldehyde treatments. When compared with the bead-milled control sample, F= 10245.62 and F=149.09, p < 0.0001 were observed for autoclaving and microwaving, respectively (Supplementary **Table 1**). A post-hoc Tukey test showed that all the autoclaving treatments (P1, P2, P3 and P4) and control belonging group differed significantly at p < .05 (table not shown) on the PCR-ability of release DNA from *E. coli* cultures. Regarding microwaving, the Tukey test showed significant (p < .05) differences from 60 to 100 s exposure times.

**Figure 4.**
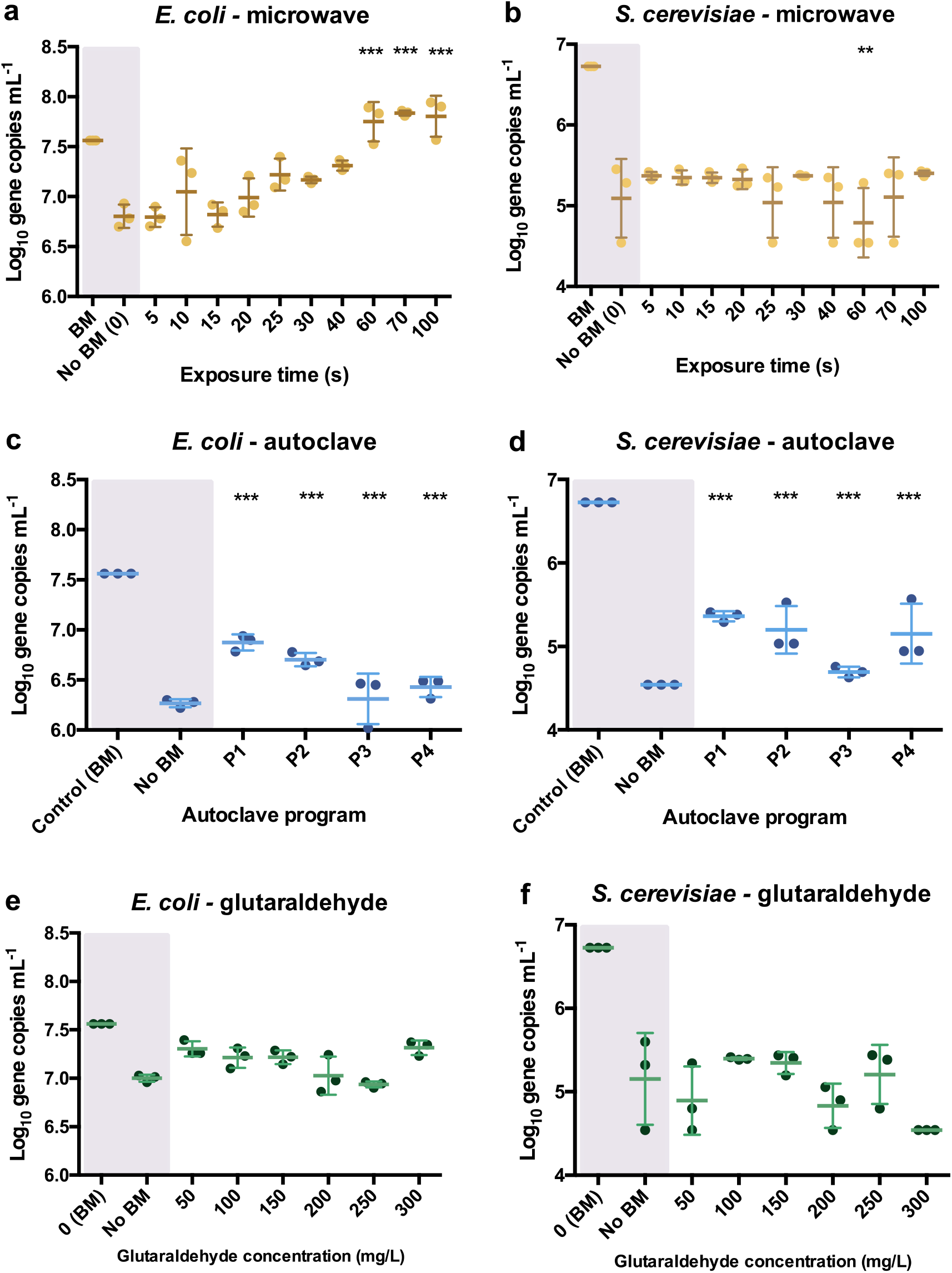
qPCR results after the different sterilization methods. Number of DNA copies obtained on supernatant of pure *E. coli* and *S. cerevisiae* cultures after microwave (A-B), autoclave (C-D) and glutaraldehyde (E-F) treatments. The results are expressed in log_10_ copies per mL. Autoclave programs: P1 (110 °C – 20 min), P2 (110 °C – 30 min), P3 (121 °C – 20 min) and P4 (121 °C – 30 min). BM: Bead-Mill treatment. ANOVA followed by post-hoc Tukey test, (**) p<0.05, (***) p<0.005.

In *S. cerevisiae* cultures, DNA was released in a constant trend when microwaving meaning that lower amounts of DNA were available in the supernatant per time unit. This balance of DNA release seems to be proportional to degradation over time. This was reflected in Fig. 4**b**, where constant values around 1.5 log_10_ differences are observed over time. A decrease of 4.78 ± 0.18 log_10_ DNA copies per mL at 60 s was observed but went up to 5.11 ± 0.2 log_10_ DNA copies per mL at 70 s. After 100 s, 5.4 ± 0.02 log_10_ DNA copies increased indicating poor loss of PCR ability even when long microwave exposure times were applied. DNA release pattern and PCR ability from *S. cerevisiae* differ drastically from the exponential released observed in *E. coli* cultures (Fig. 4**a**). After autoclaving, qPCR still detected some sequences (1.62 ± 0.15 log_10_ difference) even after the most intensive programs (P4: 121°C, 30 min, Fig. 4**d**). However, P3 (121°C, 20 min) resulted in lower amplifiable DNA sequences (2.08 ± 0.03 log_10_ difference).

As observed in *E. coli* cultures, glutaraldehyde did not have a significant effect on DNA PCR ability mainly because samples released non-detectable levels of DNA (Fig. 4**f**). An ANOVA on these scores yielded significant variation among autoclaving treatment but not after microwave and glutaraldehyde treatments. When compared with the bead-milled control sample, F=137.77, p < 0.0001 were observed for autoclaving (Supplementary **Table 2**). Tukey test showed that all the autoclaving treatments (P1, P2, P3 and P4) and control belonging group differed significantly at p < .05 (table not shown) on the PCR-ability of release DNA from *S. cerevisiae* cultures. Regarding microwaving and glutaraldehyde, no significant (p > .05) mean differences were observed supporting the low effect of these methods on PCR ability. High temperature in combination with high pressure massively degrades DNA even in a short period of time of 5 min ^18^. Some studies have shown that dry autoclaving at 100 °C for 10 min has been already sufficient to result in not amplifiable DNA from soybean ^6^. However, genomic DNA amplification has been observed at different autoclaving times (from 10 to 40 min) out of cultures of *Pseudomonas aeruginosa, Salmonella Nottingham*, and *Escherichia coli*. Collectively, this suggests a potential risk made by residual genomic DNA after inactivation of microbial cells due to potential horizontal gene transfer phenomena ^35^. Little is known on the amplification capacity of free DNA after autoclaving treatment of industrial model organisms, being a source of xenogenic contamination out of industries and laboratories.

### Quality test experiments using autoclave and microwave treatments showed faster pure DNA degradation patterns when compared to pure cultures

The different sterilization treatments were tested out *in vitro* on pure phage λ DNA and analyzed by gel electrophoresis to assess afterwards their effect on free extracellular DNA.

All the autoclaving programs applied on pure λ DNA reflected neither visible bands nor smears on gel electrophoreses, meaning that complete degradations of naked λ DNA occurred (Fig. 5**a**). DNA was neither visible on gels nor available for SYBR-Gold staining. In contrast with the DNA treated by autoclaving *in vivo* (Fig. 5A-C), where intracellular DNA migrated slower and displayed higher fragment length than released DNA. The disappearing of λ DNA treated with high temperature and pressure manifested the protection of DNA by cells against damage. Slight increased pressure and temperatures decay the primary structure of double-strand DNA by hydrolyzing its chemical bonds ^36^. This significantly affects the DNA stability and causes DNA fragmentation. This is remarkable with free DNA and was confirmed by the autoclaving of pure free λ DNA, where gels did exhibit severe DNA fragmentation.

**Figure 5.**
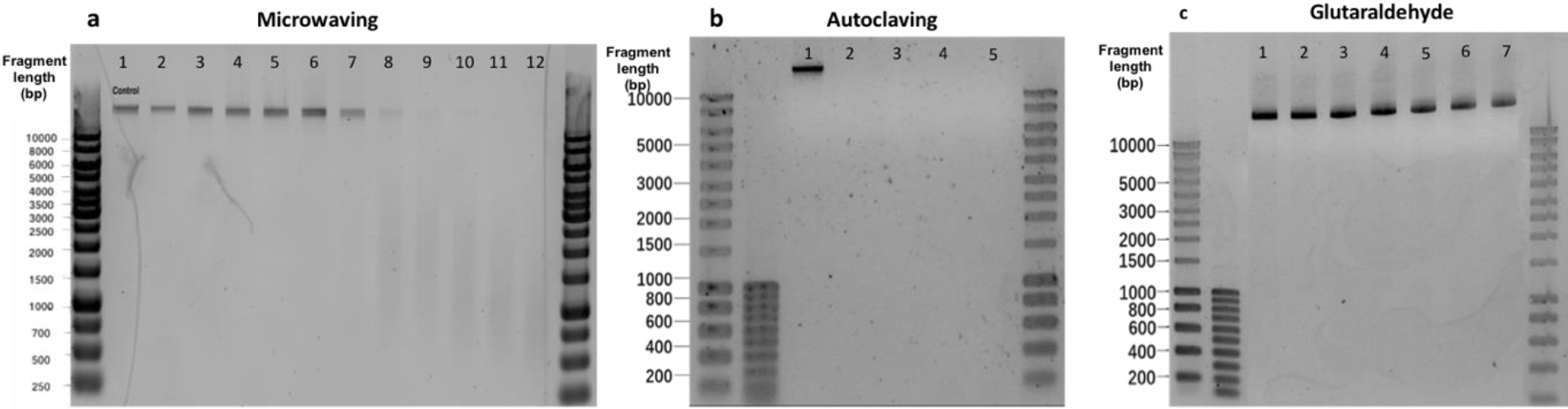
Effect of different sterilization methods on λ DNA fragmentation. (**a**) Microwaving. Lanes 1-12 correspond to 0, 5, 10, 15, 20, 25, 30, 40, 50, 60, 70, 100 s microwave exposure time, respectively. (**b**) Autoclaving. Lanes 1-5 correspond to control, P1, P2, P3 and P4, respectively. Autoclave programs: P1 (110 °C – 20 min), P2 (110 °C – 30 min), P3 (121 °C – 20 min) and P4 (121 °C – 30 min). (**c**) Glutaraldehyde. Lanes 1-7 correspond to 0, 50, 100, 150, 200, 250 and 300 mg L^−1^ glutaraldehyde, respectively.

An exposure of 30 s to microwaves was not sufficient to cause degradation and fragmentation to free pure λ DNA: the band obtained after treatment (Fig. 5**c**. Lane 7) remained identical to the untreated template (Fig. 5**c**. Lane 1). At 40 s, the DNA band lost its intensity, resulting in a smear with lower fragment lengths. After 60 s, the band disappeared and completely turned into a smear. In comparison with the results in Fig. 4, the phage λ DNA treated with the same duration of microwaving resulted in slower migrations across the gel than the DNA released from *E. coli* and *S. cerevisiae* cultures. Non-thermal factors are all of the effects that are not caused by an increase of temperature, especially when low frequencies and intensities are applied ^37^. Belyaev (2005) has shown that radiation-induced DNA breaks could not be repaired after non-thermal microwave exposure. This effect inhibits DNA repair being a plausible cause for cells inactivation. When moving from the cellular to the DNA level, microwaves can destroy DNA by denaturation, degradation, and fragmentation ^10,39^.

No DNA degradation nor fragmentation could be observed when increasing glutaraldehyde concentrations (Fig. 5**c**).

### Pure λ DNA amplification efficiency decreased under long microwave exposures and all autoclaving programs

The qPCR of the pure bacteriophage λ *int* gene was used to evaluate the extracellular DNA capacity to be amplified right after the different sterilization methods.

All the autoclave treatments were shown to significantly affect the PCR ability of the λ DNA (Fig. 6**b**) notably when applying program P4 (121 ^<u>°</u>^C, 30 min) when compared with the non-treated λ DNA control. Differences of 2.56 ± 0.61, 3.04 ± 1.22, 4.75 ± 0.24, 4.96 ± 0.28 logs were observed for P1, P2, P3 and P4 versus the control, respectively. An ANOVA on these scores yielded significant variation among autoclaving treatments when compared with the control, F= 11.41, p < 0.001 (Supplementary **Table 3**). A Tukey test showed that all the autoclaving treatments (P1, P2, P3 and P4) and control belonging group differed significantly at p < .05 (table not shown) on the PCR-ability of λ DNA.

**Figure 6.**
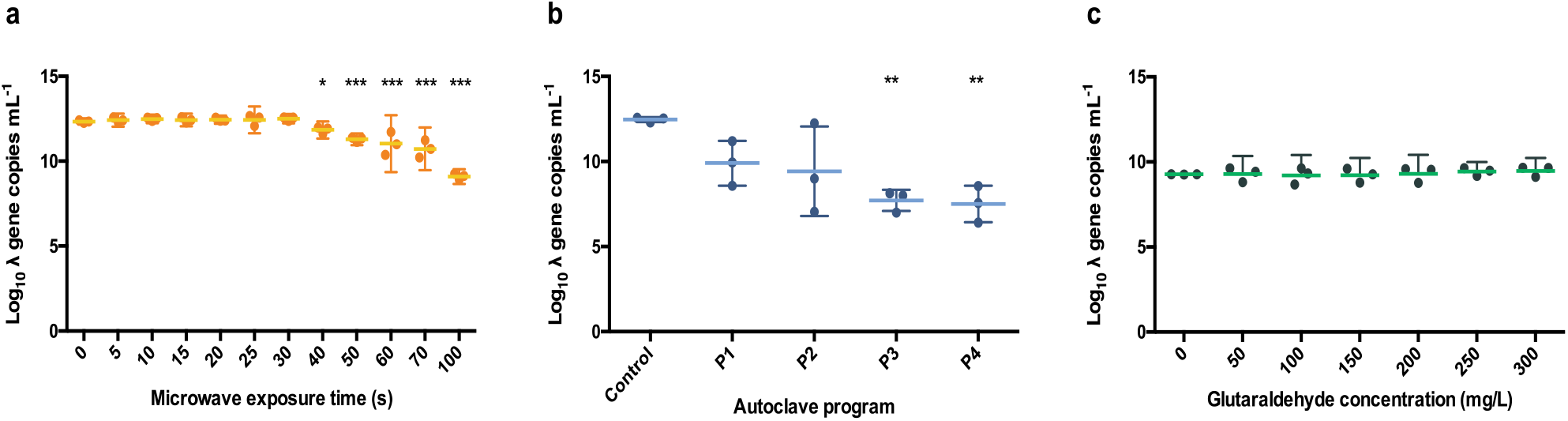
Number of *int* copies from λ DNA available after microwaving (**a**), autoclaving (**b**) and treating with glutaraldehyde (**c**). The results are expressed in log_10_ copies per mL. The middle line represents the mean, and the whiskers represent the 95% CI. Autoclave programs: P1 (110 °C – 20 min), P2 (110 °C – 30 min), P3 (121 °C – 20 min) and P4 (121 °C – 30 min). Control: non-sterilized samples corresponding to 1 g L^−1^ λ DNA.

Apart from the genome size differences between the DNAs of *E. coli* (approximately 4 Mb) and *S. cerevisiae* (approximately 12.1 Mb), which are both larger than λ DNA (48 kb) ^40,41^, the degree of DNA damage was less severe in pure λ DNA than in released DNA from microbial cells under microwave treatment. When cells are treated with microwaves, temperature in cell suspensions with higher cell concentrations increased faster than with low cell concentration ^24^. Hence, suspensions of *E. coli* and *S. cerevisiae* could end up with a higher temperature compared to the pure λ DNA solution when treated with the same duration of microwave, which accelerates the damage to the DNA in cell suspension medium and cell structures. Even if DNA in suspension could end up with higher temperatures, first it should be released.

It took around 30 to 50 s to detect high concentrations of DNA in the extracellular fraction and around more than 100 s to start seeing a decay in PCR ability. From the combination of experiments with pure cultures and free-floating λ DNA, exposure time longer than 100 s would be necessary to inactivate cells, release DNA, and fragment it. Otherwise, no effect will be observed on DNA integrity.

Similar results were observed when samples were exposed for over 40 s, 0.5 ± 0.09 logs difference, under microwaves (Fig. 6**a**). Significant variation among autoclaving treatments when compared with the control, F= 26.04, p < .0001 (Supplementary Table 4) was observed. Moreover, a Tukey test showed that exposure times over 60 s, 1.31 ± 0.17 logs difference, differed significantly at p < 0.05 (table not shown). This is supporting the clear band loss after 60 s during gel electrophoresis (Fig. 5**c**). At 100 s, a difference of 3.23 ± 0.06 logs with the untreated control sample was observed.

In contrast, the glutaraldehyde treatments did not affect the amplification capacity of pure λ DNA (Fig. 6**c**). An ANOVA on these scores did not yield significant variation among the different concentrations of glutaraldehyde when compared with the control (F= 0.70, p > 0.658, Supplementary Table 4), supporting the results obtained from the DNA fragmentation experiments (Fig. 5**b**). No significant effect on amplification ability was observed when a standard incubation time of 20 min was applied. Glutaraldehyde damages DNA ^42^ and compromises the PCR ability of DNA after some days of incubation only ^43^. In clinical procedures, an exposure of 20 min at 20 °C in a 2% w/w glutaraldehyde solution is solely used to disinfect medical instruments ^44^. We showed that this short incubation time of 20 min was not enough to impact the integrity and PCR ability of DNA. Overall, glutaraldehyde offers an efficient way to disinfect and contain DNA and xenogenic elements inside cells.

### General remarks, alternatives and perspectives

Overall, all common sterilization methods here tested are effective to inactivate microorganisms, highlighting short incubation time of 20 min with glutaraldehyde for its capacity to avoid DNA release. In terms of DNA loss of integrity, autoclave is shown to be the most effective method. However, integrity of released DNA is not completely compromised as shown by qPCR results. This opens a window for improvement in case total extracellular DNA degradation was desired. Alternatives to standard procedures are combination of methods here tested or further steps towards total removal such as ethylene oxide that has been shown to reduce DNA amplification when long exposure times are applied ^45^. Fragmented sequences as short as 20 bp haven been shown to be taken up and incorporated into the bacterial DNA, including mammoth DNA ^46^. Even if the majority of short residual DNA fragments will be re-metabolized in case they are taken up, there is a probability to be genome integrated generating new diversity ^47^.

Horizontal gene transfer phenomena from sterilized cultures may exchange all kind of DNA fragments as soon as these enter microbiome hotspots such as wastewater treatment plants ^3^. New microbial diversity can be generated through gene transfer, but also undesirable fragments such as ARGs and pathogenic islands could be unfavorably mobilized by microorganisms ^48,49^. This underlies the ubiquity and potentiality of these DNA fragments generated after sterilizations. Further research on the quality and composition of released DNA as well as rates of horizontal gene transfer are necessary to develop risk assessments strategies and to address the impact of the standard sterilization methods on biosafety and environmental and public health.

## Material and Methods

### Bacterial and yeast strains and culture preparations

A frozen stock of *Escherichia coli*, DH5α (Cell System Engineering Section, Department of Biotechnology, TU Delft, the Netherlands) was thawed and inoculated into a 300-mL flask containing 200 mL of Luria-Bertani (LB) cultivation broth. This culture was incubated in a rotary shaker for 6 h at 37 °C, 200 rpm, where late log phase was reached. A frozen stock of *Saccharomyces cerevisiae*, CEN.PK (Cell System Engineering Section, Department of Biotechnology, TU Delft, the Netherlands) was thawed and inoculated into a 500-mL flask containing 200 mL of yeast extract-peptone-dextrose (YEPD) broth composed of 1% yeast extract, 2% peptone, and 2% glucose/dextrose. This culture was incubated in a rotary shaker for 6 h at 30 °C and 200 rpm. Samples of 5 mL at 10^11^ CFU L^−1^ of cell cultures were prepared for further sterilization treatments.

### Physical sterilization treatment by microwaving

A microwave oven (Bestron Model ER-M18, 2450 MHz, 230V, 850W) with a rotating table was used. A sealed glass bottle containing 5 mL of each microorganism was placed at the center of the rotating table, 20 cm away from the irradiation source. The microwave was irradiated for a maximum of 30 s within which different time intervals were taken. For each interval time point (0, 5, 10, 12, 14, 16, 18, 20, 25 and 30 s), different sampling tubes were used to ensure that the treatment duration was continuous. Microwave was set at maximum power mode in order to avoid its automatic on and off switching. Quality controls were performed with pure 1 ng λ DNA μL^−1^ (Thermo Fisher Scientific Inc., Waltham, MA, USA) by irradiating the sample with microwaves for a maximum of 100 s.

### Chemical sterilization treatment by glutaraldehyde

A generic buffer was prepared by mixing 10 g of sodium bicarbonate (NaHCO_3_) with 90 g of disodium hydrogen phosphate (Na_2_HPO_4_). This buffer mixture was used to adjust the pH of the glutaraldehyde (C_5_H_8_O_2_) solution to an alkaline value of 8.0 which ensures the bactericidal activity of glutaraldehyde ^20^. A 2% (w/w) glutaraldehyde solution (20 g/L, 0.2 M) was prepared and the final pH of the mixture was set at 8.0. Volumes of 12 mL of microorganism suspensions were collected into 15-mL Falcon tubes and were centrifuged at 6000 x g at 4°C for 15 min. Afterwards, the pellets were resuspended in 12 mL of 1x PBS solution (pH 7.4) and placed in an 18°C water bath for chemical sterilization. Samples were treated with final glutaraldehyde concentrations of 50, 100, 150, 200, 250, and 300 mg L^−1^. Each cell suspension was tested out for 20 min. After reaction, cells were washed and resuspended in 1xPBS solution. The same procedure was followed for quality controls performed with pure λ DNA at 1 ng μL^−1^.

### Thermal sterilization treatment by autoclaving

Sterilization by autoclaving was tested in an autoclave (SHP Steriltechnik AG, Germany) at 110 °C and 121°C and 1.1 atm. overpressure. The 5 mL cell suspensions were placed in a 25-mL glass tube inside the autoclave and subjected to sterilization under four different autoclaving default programs (program P1: 110 °C, 20 min; P2: 110 °C, 30 min; P3: 121 °C, 20 min; P4: 121 °C, 30 min). Same procedure was followed for quality controls with pure λ DNA at 1 ng μL^−1^.

All sterilization experiments were done in technical triplicates by treating three individual samples taken from each culture.

### Cell survival

After sterilization, samples of *E. coli* were plated on LB agar plates, and samples of *S. cerevisiae* were plated on YEPD agar plates (100 μL). *E. coli* cells were incubated at 37°C for 24 h, and *S. cerevisiae* cells were incubated at 30°C for 24 h. Plates which contained 30 to 300 colonies were considered suitable for cell counting ^50^. Experiments were done in technical triplicates.

### DNA quantification

After sterilization, 2 mL of each cell sample was centrifuged at 10000 x g, 4 °C for 3 min. After the first centrifugation, 2 mL of supernatant was collected. To maximize DNA recovery, the residual pellet was washed with 1 mL 1xPBS solution, centrifuged again, prior collecting 1 mL of supernatant. A total of 3 mL supernatant was obtained from each sample. Supernatants and pellets were separated and stored at −80 °C pending DNA analysis. The amount of total DNA released after sterilization was measured from the supernatant by HS dsDNA Qubit assays (Qubit 3.0, Invitrogen, California, USA) according to manufacturer’s protocol.

### DNA fragmentation

DNA samples were analyzed by gel electrophoresis with agarose at 1% (w/v) (Sigma-Aldrich, Haverhill, United Kingdom) in 1xTAE buffer. DNA was post-stained using SYBR Gold solution (10000x) (Thermo Fisher Scientific, USA) mixed in 1x TAE buffer (AppliChem, Germany) at 1/10K (v/v). Gels after running were immersed into staining buffer for 40 min and were further checked with fluorescence imaging system (Syngene, UK).

### Primers selection for *E. coli* and *S. cerevisiae* and design for λ DNA

Forward and reverse primers were designed to assess the PCR ability of the released DNA fragments after the different sterilization methods used in this study. All primers were purchased from Sigma-Aldrich (Sigma-Aldrich, Haverhill, United Kingdom). The ß-glucuronidase (*uidA*) gene from *E. coli* was evaluated by using *uidA* forward (5’-TGGTAATTACCGACGAAAACGGC-3’) and *uidA* reverse (5’-ACGCGTGGTTACAGTCTTGCG-3’) primers ^51^. The TATA binding protein-associated factor (*TAF10*) gene from *S. cerevisiae* was evaluated using *TAF10* forward (5’-ATATTCCAGGATCAGGTCTTCCGTAGC-3’) and *TAF10* reverse (5’-GTAGTCTTCTCATTCTGTTGATGTTGTTGTTG-3’) primers ^52^. The λ bacteriophage genome was obtained from GenBank (Wu, 1972; Entry Number J02459.1). Primers targeting the λ integrase (*int*) gene were designed in house using SnapGene (GSL Biotech, www.snapgene.com): λ *int* forward (5’-GTTACCGGGCAACGAGTTGG-3’), λ *int* reverse (5’-ATGCCCGAGAAGATGTTGAGC-3’) primers.

### DNA extraction and Quantitative PCR (qPCR) analysis

The quantification of the λ *int* gene, *E. coli uidA* gene, and *S. cerevisiae TAF10* gene in DNA fragments potentially released after sterilization treatments were analyzed by qPCR (QTower 3, Analytica Jena, Germany). For the standard curve construction, genomic DNA from the model organisms *E. coli* and *S. cerevisiae* was isolated using NucleoSpin^®^ Soil (Macherey-Nagel) and Yeast DNA Extraction (Thermo Fisher Scientific Inc., Waltham, MA, USA) kits, respectively. Serial dilutions from 100 ng μL ^−1^ down to 10^−5^ ng μL^−1^ were used to generate the standard curve. The validation standard curve construction was performed by purchasing 0.3 μg μL^−1^ λ pure DNA (Thermo Fisher Scientific Inc., Waltham, MA, USA). Serial dilution from 1 ng μL^−1^ down to 10^−8^ ng μL^−1^ were used to generate the standard curve. The samples were tested for amplification capacity after sterilization treatments by collecting 1 mL of sterilized culture and centrifugating it at 15000 x g for 5 min. The supernatant containing released DNA was collected and used for qPCR analysis. All qPCR reactions were performed in volumes of 20 μL composed of 10 μL of IQ™ SYBR^®^ Green Supermix (Bio-Rad), 0.2 μL of each primer at 50 μM, 8.6 μL ultrapure water (Sigma-Aldrich, Haverhill, United Kingdom) and 1 μL of template DNA.

The thermal profile selected for the λ *int* gene consisted of 5 min at 95 °C hot-start polymerase activation followed by 40 cycles of DNA dissociation at 95 °C for 30 s, primers annealing at 55 °C for 30 s fragment elongation at 72 °C for 30 s, and terminated by holding at 4 °C.

The thermal profile selected for the *E. coli uidA* gene gene consisted of 5 min at 95 °C hot-start polymerase activation followed by 40 cycles of DNA dissociation at 95 °C for 30 s, primers annealing at 57 °C for 30 s, fragment elongation at 72 °C for 30 s, and terminated by holding at 4 °C.

The thermal profile selected for the *S. cerevisiae TAF10* gene consisted of 5 min at 95 °C hot-start polymerase activation followed by 40 cycles of DNA dissociation at 95 °C for 30 s, primers annealing at 55 °C for 30 s, fragment elongation at 72 °C for 30 s, and terminated by holding at 4 °C.

### Statistical analysis

Statistical analyses were performed with R 3.5.1 ^54^ and RStudio (https://www.rstudio.com/). For the analysis and determination of the most effective parameters of the sterilization methods effect on DNA amplification a two-way ANOVA test followed by a post-hoc Tukey’s honestly significant difference (HSD) test at the 0.05 probability level were performed. Figures were prepared using GraphPad Prism 6 (GraphPad Software, San Diego, CA, USA).

## Supporting information

Calderon Franco et al. - Supplementary material

## Acknowledgements

This work is part of the research project “Transmission of Antimicrobial Resistance Genes and Engineered DNA from Transgenic Biosystems in Nature” (Targetbio) funded by the programme Biotechnology & Safety (grant no. 15812) of the Applied and Engineering Sciences Division of the Dutch Research Council (NWO).

## Authors’ contributions

DGW, BA and DCF designed the study with inputs of QL and MvL. DCF and QL performed the experimental investigations. DCF and DGW wrote the manuscript with direct contribution, edits, and critical feedback by all authors.

## Supplementary information

**Table S1.** *E. coli* and *S. cerevisiae* cell viability after different autoclaving programs. The percentage of viable cells is calculated against corresponding control sample cells (non-autoclaved cells).

**Table S2.** ANOVA results using the different methods Log_10_ *uidA* gene copies mL^−1^ after sterilization methods on *E. coli* as the criterion

**Table S3.** ANOVA results using the different methods Log_10_ *TAF10* gene copies mL^−1^ after sterilization methods on *S. cerevisiae* as the criterion

**Table S4.** ANOVA results using the different methods Log_10_ *intI* gene copies mL^−1^ after sterilization methods on λ as the criterion

**Figure S1.** Microscopic pictures of *E. coli* cells treated with microwave

**Figure S2.** Microscopic pictures of *S. cerevisiae* cells treated with microwave

**Figure S3.** Microscopic pictures of *E. coli* cells treated with autoclaving

**Figure S4.** Microscopic pictures of *S. cerevisiae* cells treated with autoclaving

**Figure S5.** Microscopic pictures of *E. coli* cells treated with glutaraldehyde

**Figure S6.** Microscopic pictures of *S. cerevisiae* cells treated with glutaraldehyde

**Figure S7.** DNA quantification (ng μL^−1^) and total DNA released (%) on the amount of DNA released after microwaving and glutaraldehyde treatment from *E. coli* and *S. cerevisiae* cultures.

**Figure S8.** DNA quantification (ng μL^−1^) and total DNA released (%) on the amount of DNA released after autoclaving from *E. coli* and *S. cerevisiae* cultures.

**Figure S9.** Electrophoretic gel of E. coli intracellular and released DNA together with S. cerevisiae intracellular and released DNA with increasing microwave exposure times.

**Figure S10.** Electrophoretic gel of E. coli intracellular and released DNA together with S. cerevisiae intracellular and released DNA with different types of autoclaving.

**Figure S11.** Electrophoretic gel of E. coli intracellular and released DNA together with S. cerevisiae intracellular and released DNA with increasing concentration of glutaraldehyde.

