## Supplementary material for "Anticipating xenogenic pollution at the source: Impact of sterilizations on DNA release from microbial cultures": Calderon Franco et al. - Supplementary material

Deposited as pre-print on *bioRxiv*

### Supplementary material

**Table S1.** *E. coli* and *S. cerevisiae* cell viability after different autoclaving programs. The percentage of viable cells is calculated against corresponding control sample cells (non-autoclaved cells).

| Treatment condition | *E. coli* | | *S. cerevisiae* | |
|---|---|---|---|---|
| | Cell number Log (CFU) $mL^{-1}$ | Viability (%) | Cell number Log (CFU) $mL^{-1}$ | Viability (%) |
| **Autoclaving program** | | | | |
| 110 °C, 20 min | *NA | *NA | *NA | *NA |
| 110 °C, 30 min | *NA | *NA | *NA | *NA |
| 121 °C, 20 min | *NA | *NA | *NA | *NA |
| 121 °C, 30 min | *NA | *NA | *NA | *NA |

*NA (not applicable): Cell concentration reduced under the accurate detection limit of 100 CFU $mL^{-1}$.

**Table S2.** Fixed-Effects ANOVA results using the different methods $Log_{10}$ *uidA* copies $mL^{-1}$ after sterilization methods on *E. coli* as the criterion

| Treatment condition | | | ***Microwaving*** | | | | |
|---|---|---|---|---|---|---|---|
| Predictor | Sum of Squares | df | Mean Square | F | p | partial η2 | partial η2 90% CI [LL, UL] |
| (Intercept) | 149.09 | 1 | 149.09 | 4795.50 | .000 | | |
| Exposure time (s) | 5.33 | 11 | 0.48 | 15.57 | **.000** *** | .88 | [.69 .88] |
| Error | 0.75 | 24 | 0.03 | | | | |

| Treatment condition | | | ***Autoclaving*** | | | | |
|---|---|---|---|---|---|---|---|
| Predictor | Sum of Squares | df | Mean Square | F | p | partial η2 | partial η2 90% CI [LL, UL] |
| (Intercept) | 172.84 | 1 | 172.84 | 10246.62 | .000 | | |
| Autoclave program | 3.04 | 4 | 0.76 | 45.10 | **.000** *** | .95 | [.81, .96] |
| Error | 0.17 | 10 | 0.02 | | | | |

| Treatment condition | | | ***Glutaraldehyde*** | | | | |
|---|---|---|---|---|---|---|---|
| Predictor | Sum of Squares | df | Mean Square | F | p | partial η2 | partial η2 90% CI [LL, UL] |
| (Intercept) | 156.07 | 1 | 156.07 | 6266.22 | 0.000 | | |
| Glutaraldehyde concentration (mg $L^{-1}$) | 0.23 | 6 | 0.04 | 1.53 | .240 | .40 | [.00, .46] |
| Error | 0.35 | 14 | 0.02 | | | | |

Note. LL and UL represent the lower-limit and upper-limit of the partial η2 confidence interval, respectively.

**Table S3.** Fixed-Effects ANOVA results using the different methods $Log_{10}$ *TAF10* copies $mL^{-1}$ after sterilization methods on *S. cerevisia*e as the criterion

| Treatment condition | | | ***Microwaving*** | | | | |
|---|---|---|---|---|---|---|---|
| Predictor | Sum of Squares | df | Mean Square | F | p | partial η2 | partial η2 90% CI [LL, UL] |
| (Intercept) | 85.85 | 1 | 85.85 | 960.59 | .000 | | |
| Exposure time (s) | 3.88 | 11 | 0.35 | 3.95 | **.002** ** | .64 | [.21, .65] |
| Error | 2.15 | 24 | 0.09 | | | | |

| Treatment condition | | | | ***Autoclaving*** | | | |
|---|---|---|---|---|---|---|---|
| Predictor | Sum of Squares | df | Mean Square | F | p | partial η2 | partial η2 90% CI [LL, UL] |
| (Intercept) | 137.77 | 1 | 137.77 | 2660.87 | .000 | | |
| Autoclave program | 7.48 | 4 | 1.87 | 36.10 | **.000** *** | .94 | [.77, .95] |
| Error | 0.52 | 10 | 0.05 | | | | |

| Treatment condition | | | | ***Glutaraldehyde*** | | | |
|---|---|---|---|---|---|---|---|
| Predictor | Sum of Squares | df | Mean Square | F | p | partial η2 | partial η2 90% CI [LL, UL] |
| (Intercept) | 87.39 | 1 | 87.39 | 666.34 | .000 | | |
| Glutaraldehyde concentration (mg $L^{-1}$) | 0.81 | 6 | 0.14 | 1.04 | .447 | .31 | [.00, .36] |
| Error | 1.84 | 14 | 0.13 | | | | |

Note. LL and UL represent the lower-limit and upper-limit of the partial η2 confidence interval, respectively.

**Table S4.** Fixed-Effects ANOVA results using the different methods $Log_{10}$ λ *int* gene copies $mL^{-1}$ after sterilization methods as the criterion

| Treatment condition | | | | ***Autoclaving*** | | | |
|---|---|---|---|---|---|---|---|
| Predictor | Sum of Squares | df | Mean Square | F | p | partial η2 | partial η2 90% CI [LL, UL] |
| (Intercept) | 466.00 | 1 | 466.00 | 228.31 | 0.000 | | |
| Autoclave program | 48.25 | 4 | 12.06 | 5.91 | **0.01 **** | .70 | [.17, .76] |
| Error | 20.41 | 10 | 2.04 | | | | |

| Treatment condition | | | | ***Microwaving*** | | | |
|---|---|---|---|---|---|---|---|
| Predictor | Sum of Squares | df | Mean Square | F | p | partial η2 | partial η2 90% CI [LL, UL] |
| (Intercept) | 456.43 | 1 | 456.43 | 5556.42 | 0.000 | | |
| Exposure time (s) | 36.32 | 11 | 3.30 | 40.20 | **.000 ***** | .95 | [.87, .95] |
| Error | 1.92 | 24 | 0.08 | | | | |

| Treatment condition | | | | ***Glutaraldehyde*** | | | |
|---|---|---|---|---|---|---|---|
| Predictor | Sum of Squares | df | Mean Square | F | p | partial η2 | partial η2 90% CI [LL, UL] |
| (Intercept) | 257.46 | 1 | 257.46 | 1925.41 | 0.000 | | |
| Glutaraldehyde concentration (mg $L^{-1}$) | 0.19 | 6 | 0.03 | 0.24 | .956 - | .0 | [.00, 1.00] |
| Error | 1.87 | 14 | 0.13 | | | | |

Note. LL and UL represent the lower-limit and upper-limit of the partial η2 confidence interval, respectively.

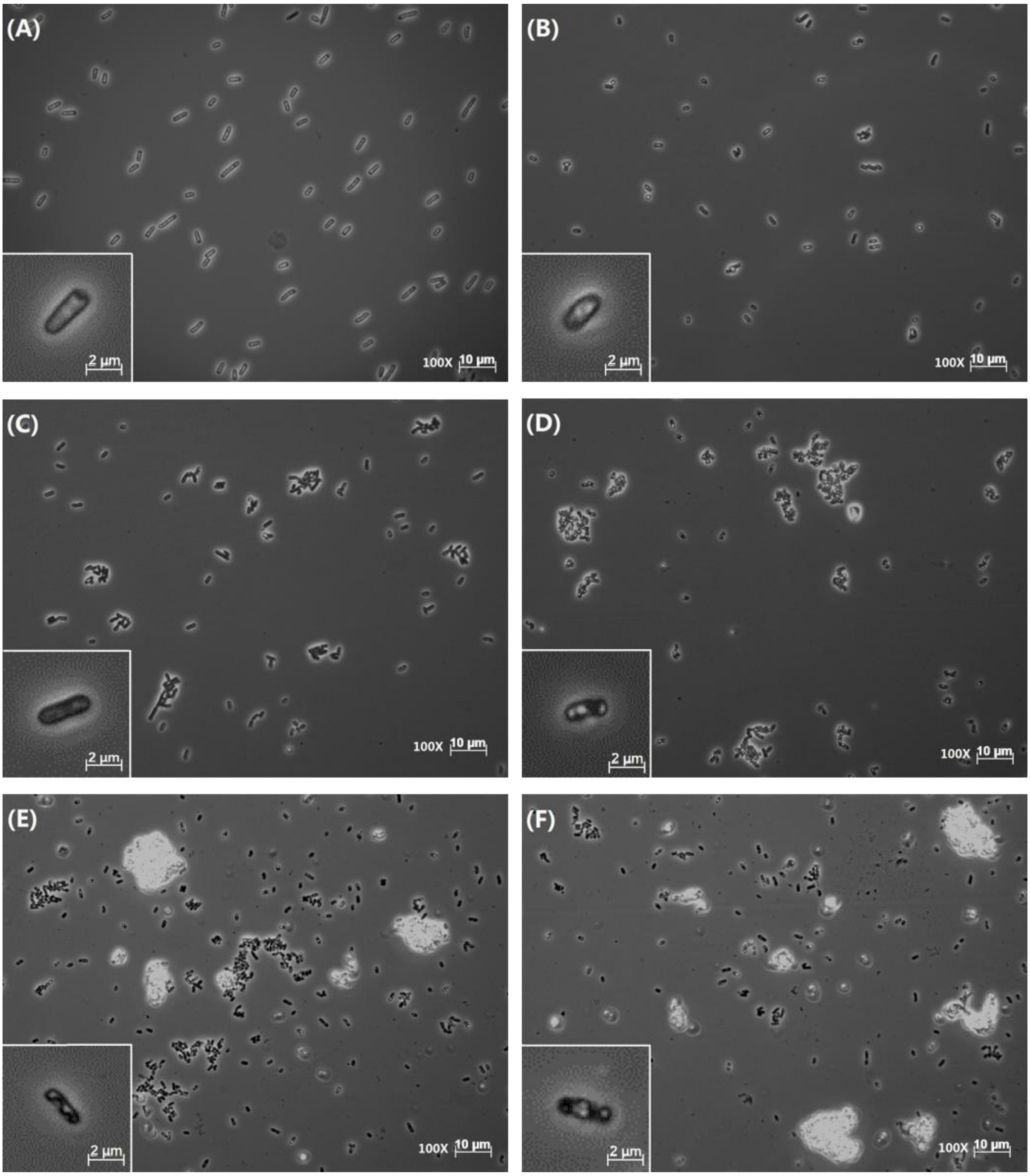


**Figure S1.** Microscopic pictures of *E. coli* cells treated with microwave (2450 MHz, 230V, 850W) sterilization method at 10 s **(b)**, 15 s **(c)**, 20 s **(d)**, 25 s **(e),** and 30 s **(f)** in comparison with 0 s untreated control cells **(a)**. The morphological structure of single cells at each time point are shown at bottom left.

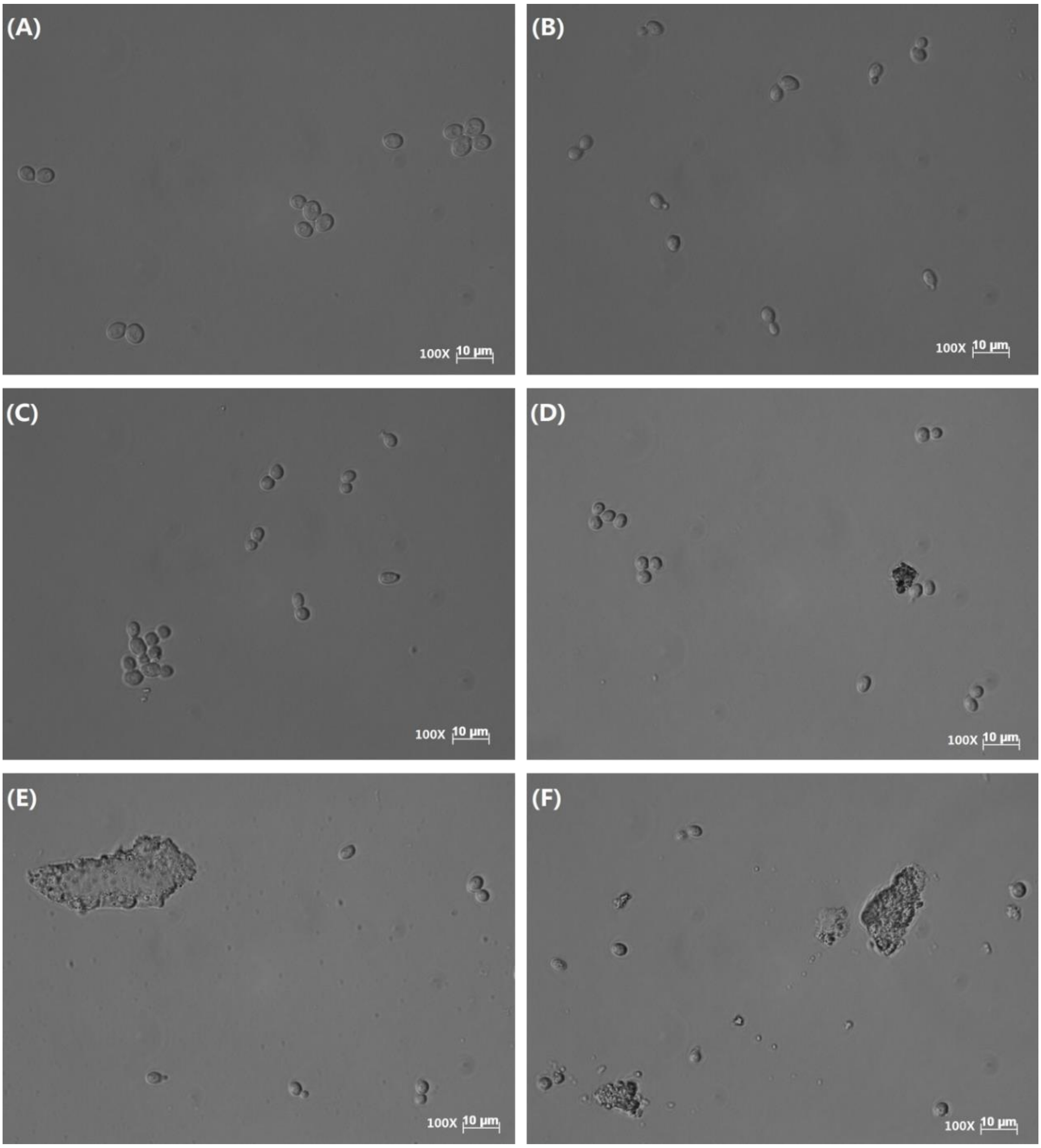


**Figure S2.** Microscopic pictures of *S. cerevisiae* cells treated with microwave (2450 MHz, 230V, 850W) sterilization method at 10 s **(b)**, 15 s **(c)**, 20 s **(d)**, 25 s **(e),** and 30 s **(f)** in comparison with 0 s untreated control cells **(a)**.

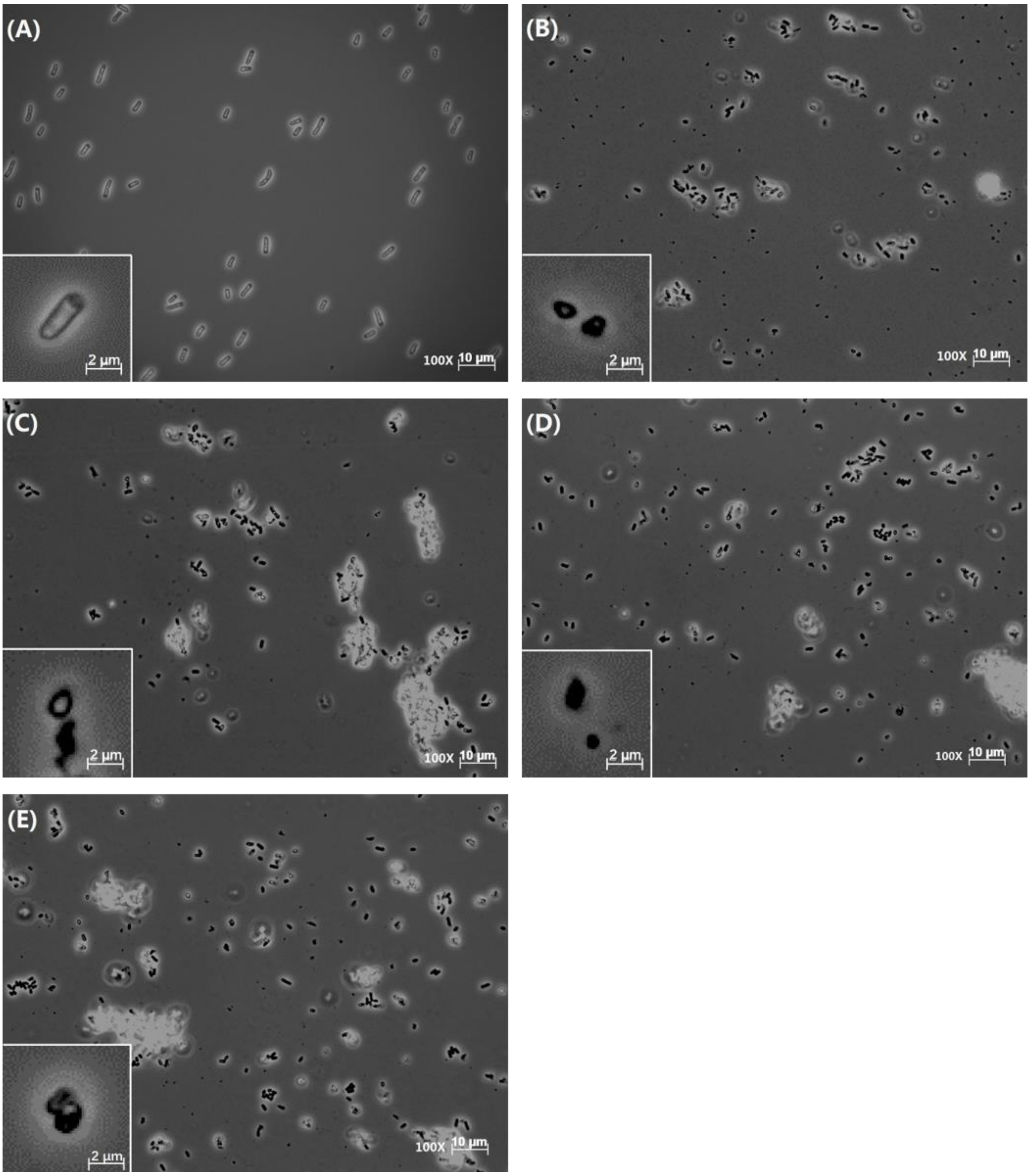


**Figure S3.** Microscopic image of *E. coli* cells through different type of autoclaving program. Cells after autoclaving program of 110 °C, 20 min (**b**), 110 °C, 30 min (**c**), 121 °C, 20 min (**d**), 121 °C, 30 min (**e**) are in comparison with untreated control cells (**a**). The morphological structure of single cells under each types of autoclaving program are shown at bottom left.

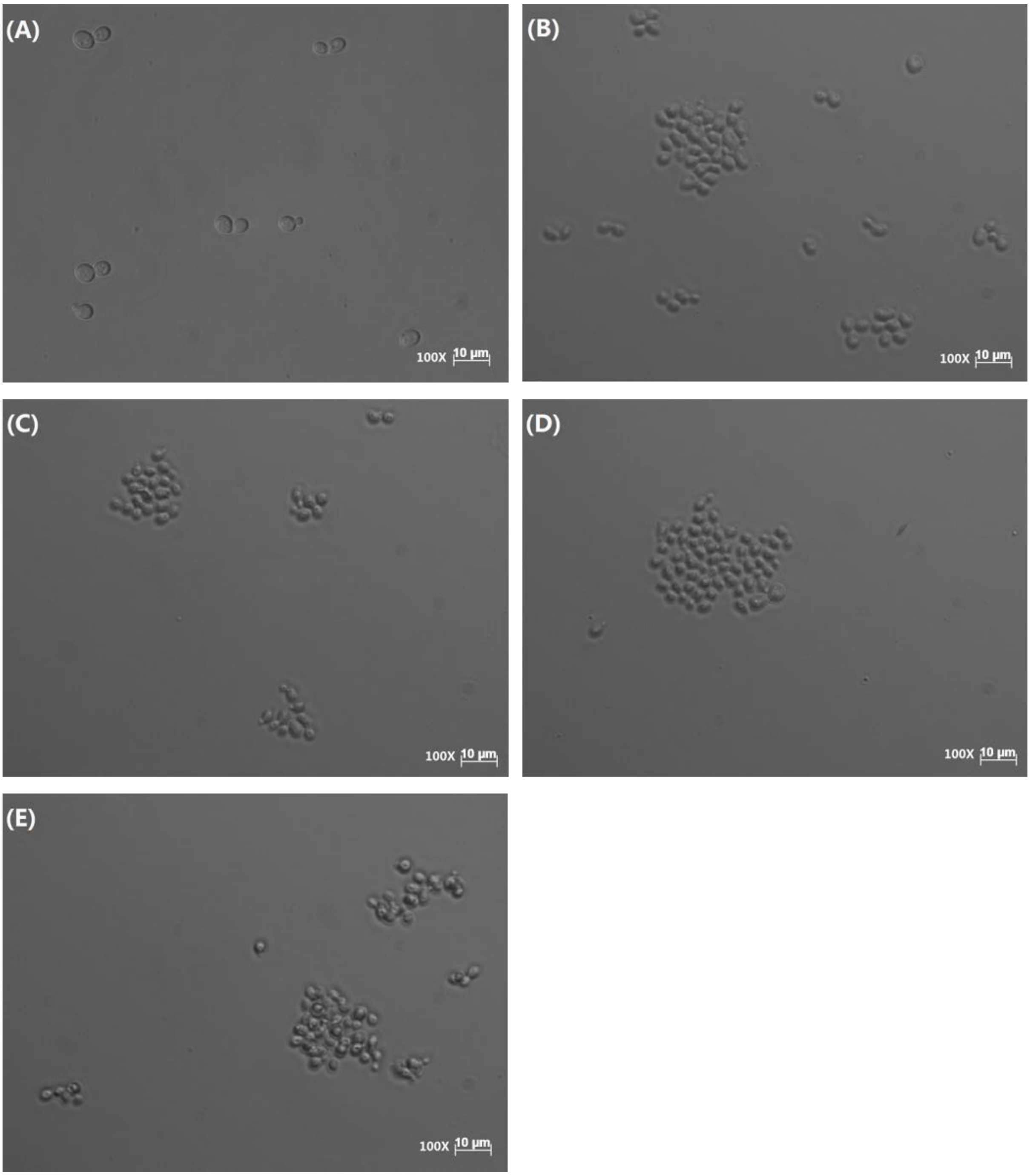


**Figure S4.** Microscopic image of *S. cerevisiae* cells through different type of autoclaving program. Cells after autoclaving program of 110 °C, 20 min (**b**), 110 °C, 30 min (**c**), 121 °C, 20 min (**d**), 121 °C, 30 min (**e**) are in comparison with untreated control cells (**a**).

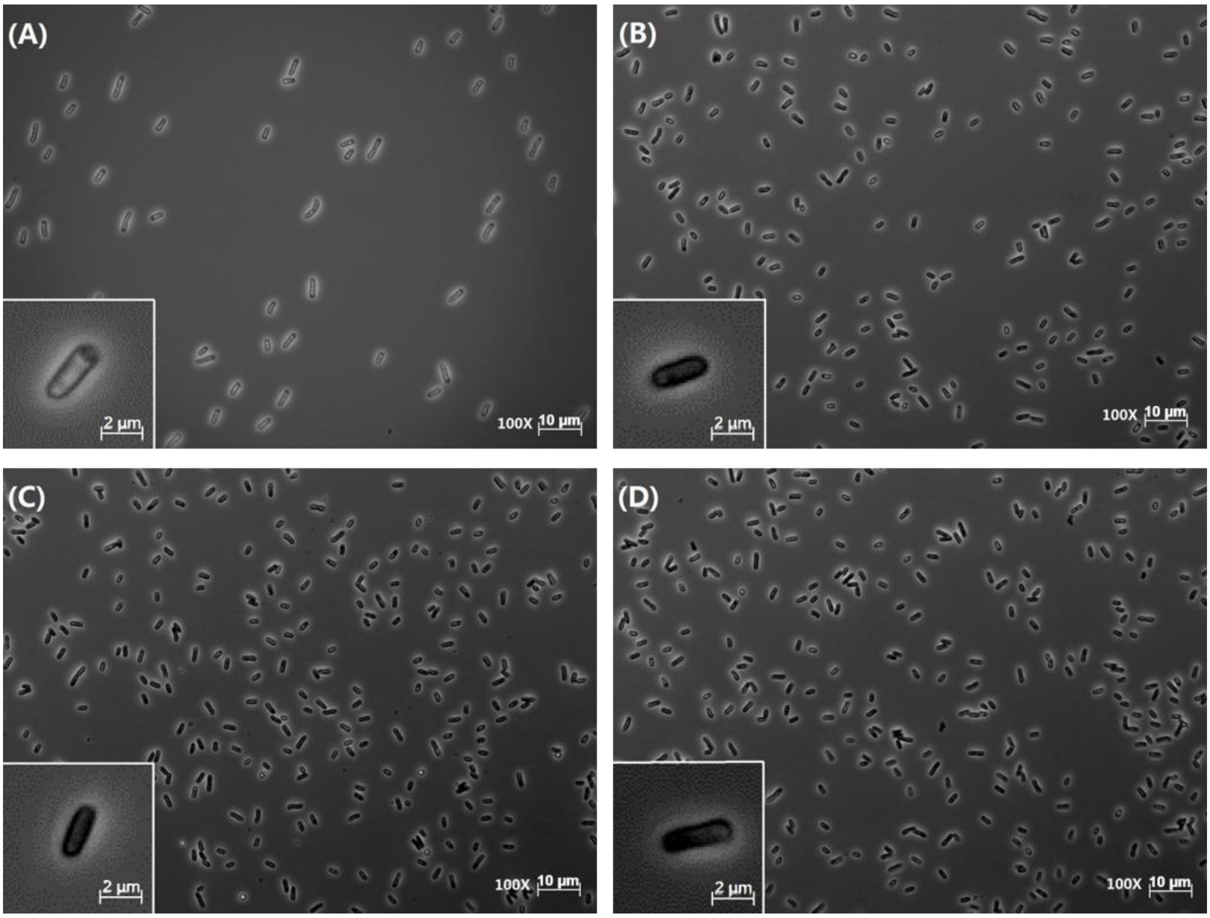


**Figure S5.** Microscopic pictures of *E. coli* cells treated with various concentrations of glutaraldehyde after 20 min incubation time. Cells affected by 100 mg $L^{-1}$ (**b**), 200 mg $L^{-1}$ (**c**), and 300 mg $L^{-1}$ (**d**) of glutaraldehyde are in comparison with untreated control cells (**a**). The morphological structure of single cells treated with each dose of glutaraldehyde are shown at bottom left.

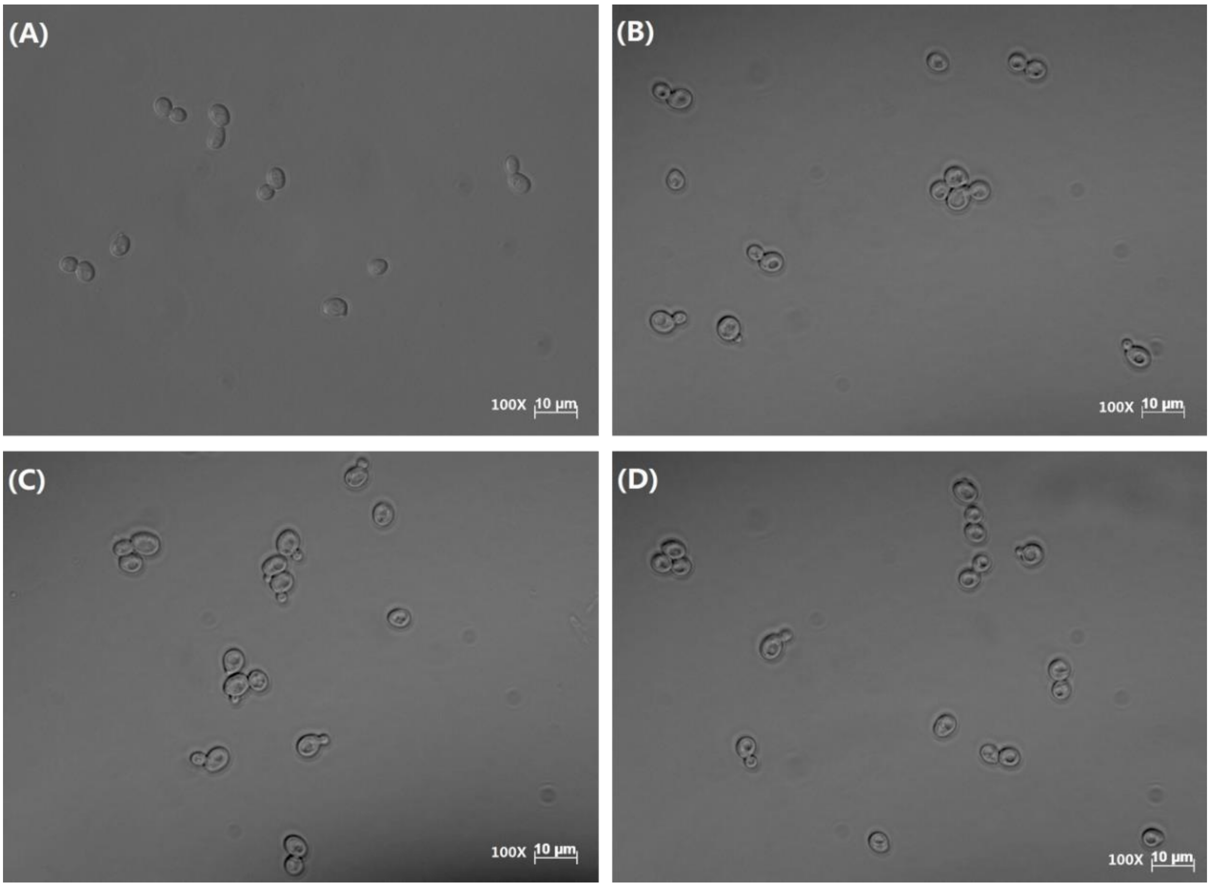


**Figure S6.** Microscopic pictures of *S. cerevisiae* cells treated with various concentrations of glutaraldehyde after 20 min incubation time. Cells affected by 100 mg $L^{-1}$ (**b**), 200 mg $L^{-1}$ (**c**), and 300 mg $L^{-1}$ (**d**) of glutaraldehyde are in comparison with untreated control cells (**a**).

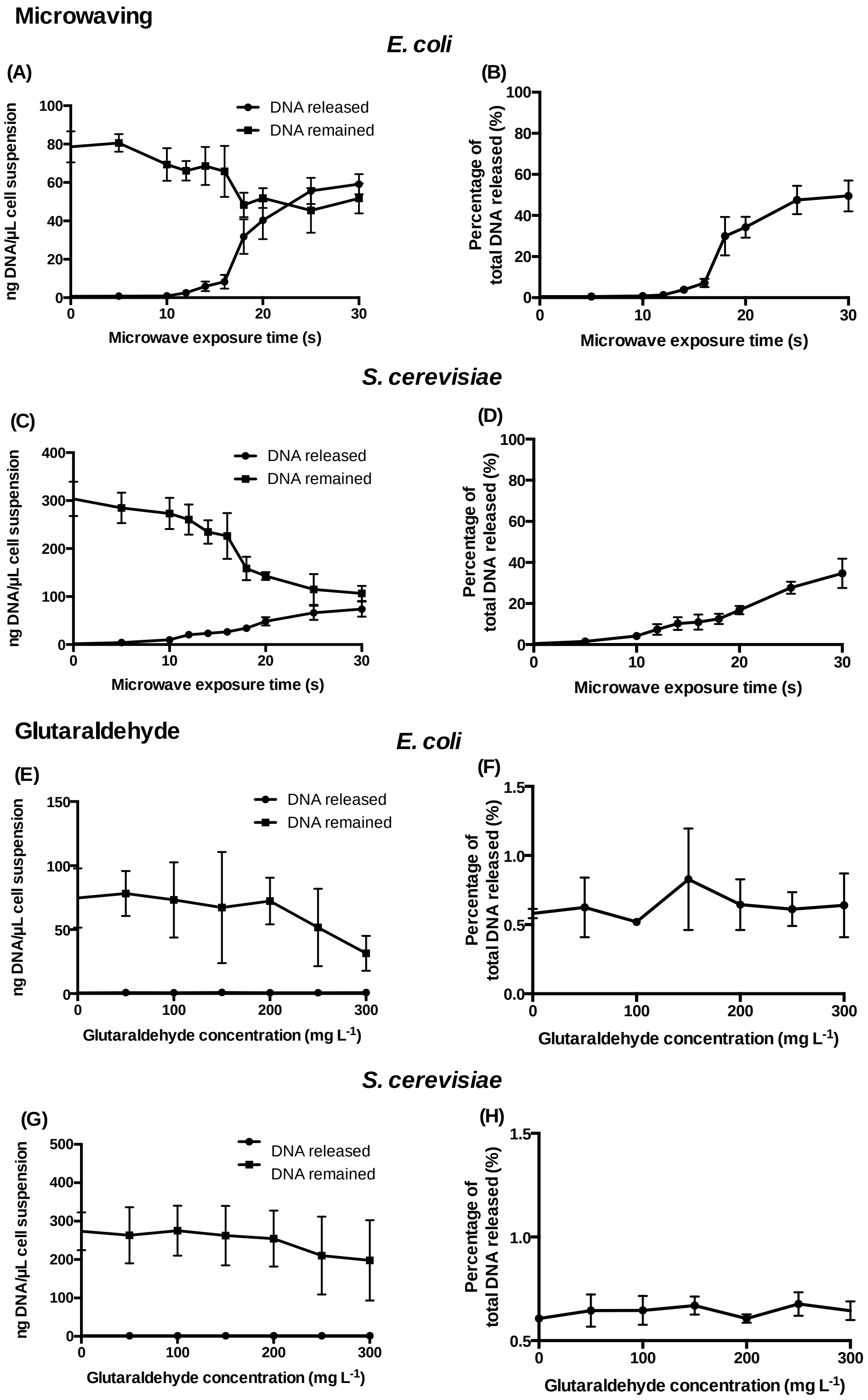


**Figure S7.** DNA quantification measurement on the amount of DNA released and remained of *E. coli* (B) and *S. cerevisiae* (C) treated with different microwave exposure times. DNA released and remained of *E. coli* (E) and S. cerevisiae (G) treated with different glutaraldehyde concentrations. Total DNA released from *E. coli* (B) and *S. cerevisiae* (D) treated with different microwave exposure times. Total DNA released from *E. coli* (F) and *S.*

*cerevisiae* (H) treated with different glutaraldehyde concentrations. The percentage shows the ratios of the amount of DNA released in the supernatant against the total amount of DNA (released and remained combined).

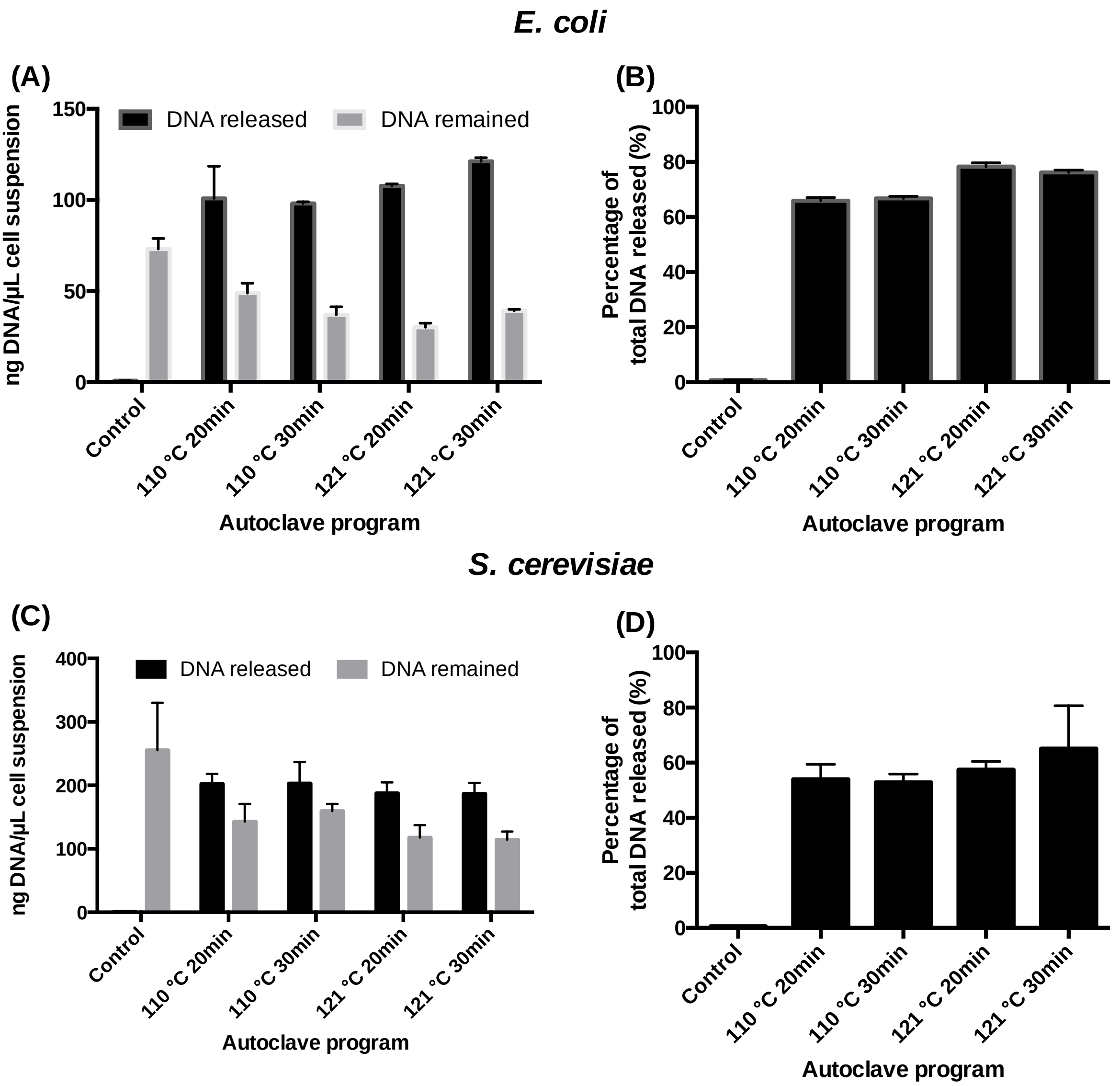


**Figure S8.** DNA quantification measurement on the amount of DNA released and remained of *E. coli* (A) and *S. cerevisiae* (C) treated with four different autoclaving programs. Total DNA released from *E. coli* (B) and *S. cerevisiae* (D) treated with four different autoclaving programs. The percentage shows the ratios of the amount of DNA released in the supernatant against the total amount of DNA.

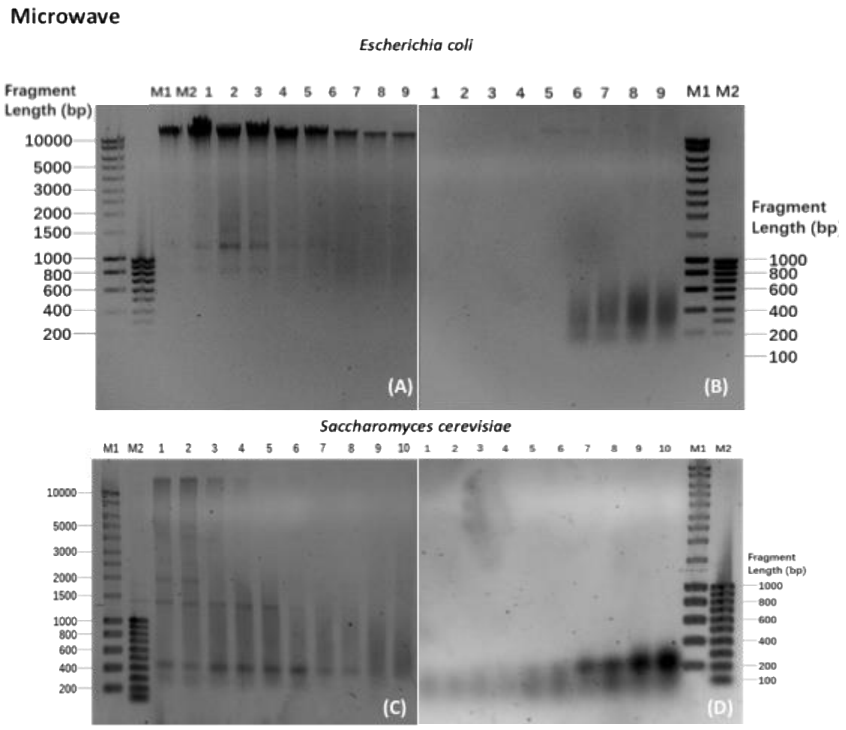


**Figure S9.** Electrophoretic gel of *E. coli* intracellular (A) and released DNA (B) together with *S. cerevisiae* intracellular (C) and released DNA (D) with increasing microwave exposure times. Lanes 1-10: *S. cerevisiae* intracellular DNA collected at 0, 10, 12, 14, 16, 18, 20, 25, 30, 40 s. Lanes 1-9 *E. coli* intracellular DNA collected at 0, 10, 12, 14, 16, 18, 20, 25, 30, 40 s.

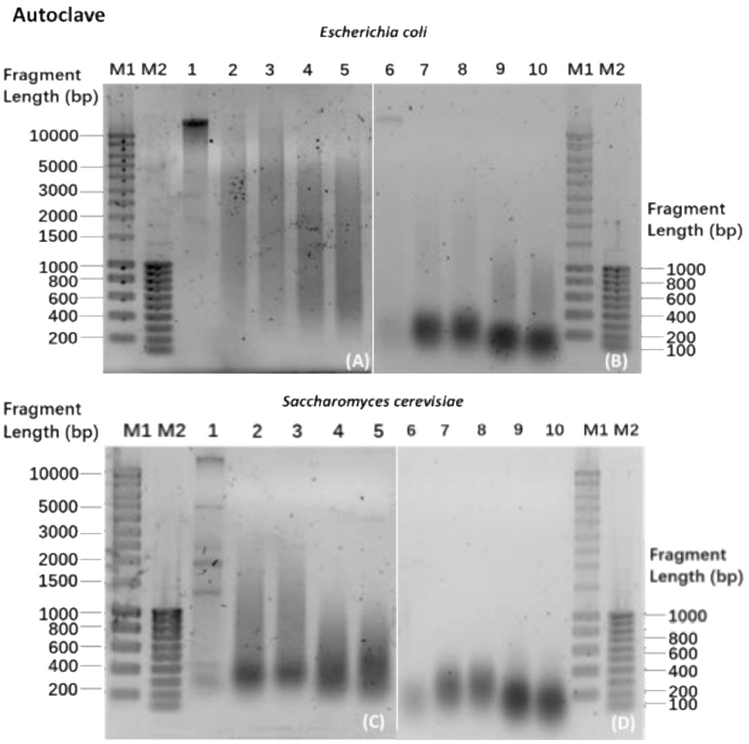


**Figure S10.** Electrophoretic gel of *E. coli* intracellular (A) and released DNA (B) together with *S. cerevisiae* intracellular (C) and released DNA (D) with different types of autoclaving. Lane 1: control sample intracellular DNA. Lane 2-5: intracellular DNA treated with P1, P2, P3 and P4. Lane 6: control sample released DNA. Lane 7-10: released DNA treated with P1, P2, P3 and P4 autoclaving programs. Autoclave programs: P1 (110 ºC – 20 min), P2 (110 ºC – 30 min), P3 (121 ºC – 20 min) and P4 (121 ºC – 30 min).

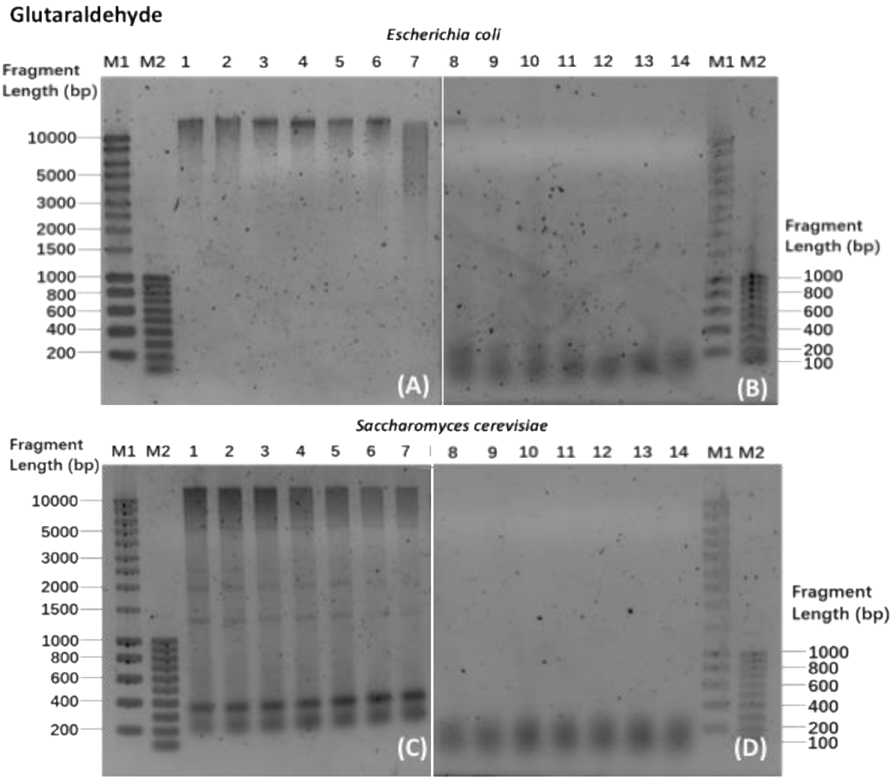


**Figure S11.** Electrophoretic gel of *E. coli* intracellular (A) and released DNA (B) together with *S. cerevisiae* intracellular (C) and released DNA (D) with increasing concentration of glutaraldehyde. Lanes 1-7: intracellular DNA collected at 0, 5, 100, 150, 150, 200, 300 mg $L^{-1}$ glutaraldehyde. Lanes 8-14: released DNA collected at 0, 50, 100, 150, 200, 300 mg $L^{-1}$ glutaraldehyde.
